# Auditory Profiles in Tinnitus are Age-Dependent: Electrophysiological and Behavioral Evidence

**DOI:** 10.64898/2026.06.02.729537

**Authors:** Pauline Devolder, Hannah Keppler, Ingeborg Dhooge, Sarah Verhulst

## Abstract

Tinnitus is commonly associated with hearing loss, yet it can also occur in individuals with clinically normal audiometric thresholds. This dissociation has led to the hypothesis that hidden sensorineural hearing loss underlies tinnitus in audiometrically normal-hearing individuals. However, identifying such subclinical deficits non-invasively is challenging because audiometric measures are influenced by age-related changes and interactions among sensorineural processes. In this study, we disentangled the contributions of tinnitus, age, and hearing status to sensorineural encoding and speech perception. We included 113 participants, divided into age-and hearing-status-matched groups with and without tinnitus, and assessed them using otoacoustic emissions, auditory evoked potentials, auditory reflex measurements, and behavioral tasks of speech perception. This design enabled a rigorous evaluation of whether hidden sensorineural deficits underlie tinnitus.

Age and hearing status had substantial effects on objective measures of sensorineural function, whereas tinnitus-related effects were subtle and age specific. Young adults with tinnitus and normal audiometric thresholds tend to exhibit enhanced auditory brainstem responses, elevated envelope following responses, and better vowel discrimination. In contrast, middle-aged adults with tinnitus showed no such enhancements and demonstrated poorer speech-in-noise performance. Correlation analyses revealed a tinnitus-related shift toward greater reliance on central auditory processing, compared with the predominantly peripheral associations observed in controls. The middle ear muscle reflex was not associated with tinnitus but was correlated with hyperacusis-related parameters.

Together, these findings suggest distinct tinnitus-related auditory profiles across the lifespan: neural enhancement and improved vowel discrimination in young adults, versus degraded sensorineural encoding and reduced speech intelligibility in middle-aged adults.

**Significance Statement:** Tinnitus affects a significant portion of the population, yet its underlying origins are still unclear. While hearing loss is a common cause, individuals with tinnitus may also have normal hearing thresholds. This suggests that subtle sensorineural damage may also play a role. This study critically investigates tinnitus-, age-, and hearing-related sensorineural encoding using non-invasive electrophysiological measures, auditory reflexes, and speech perception tasks in carefully matched participant groups. The study reveals distinct tinnitus-related auditory profiles throughout the lifespan; including enhanced sensorineural processing in young adults and degraded encoding with impaired speech perception in middle-aged adults. These findings provide critical insight into the mechanisms underlying tinnitus and offer objective markers for future research on tinnitus diagnosis and treatment.

## Introduction

Tinnitus is defined as the perception of a continuous sound in the absence of an external sound source (De Ridder et al., 2021). Various theories suggest that reduced peripheral input due to hearing loss may lead to central overactivity or neural noise, which the brain interprets as sound (Auerbach, Rodrigues, & Salvi, 2014; Johannesen & Lopez-Poveda, 2021; Knipper et al., 2013; Zeng, 2020). However, uncertainties remain regarding peripheral hearing. While hearing loss is a well-known cause of tinnitus, a substantial subgroup presents with normal audiometric thresholds (Langguth et al., 2017). Advances in auditory research have revealed that even with normal thresholds, underlying sensorineural deficits can occur and manifest as speech-intelligibility difficulties, referred to as hidden hearing loss (Kujawa & Liberman, 2009). It has therefore been suggested that tinnitus in normal-hearing individuals may reflect underlying hidden sensorineural hearing deficits (Knipper et al., 2013; Schaette & McAlpine, 2011).

This potential link between hidden sensorineural hearing loss and tinnitus has been explored using several auditory measurement techniques, but findings remain inconsistent. Some studies reported reduced otoacoustic emissions (OAEs), while others have not observed this (Serra et al., 2015). Auditory brainstem responses (ABRs) show reduced wave I amplitudes (Bramhall, Konrad-Martin, & McMillan, 2018; Bramhall et al., 2019; Möhrle et al., 2019) and enlarged wave V/I ratios (Sendesen et al., 2022; Song et al., 2018), often explained by higher central gain due to reduced peripheral input. However, such findings were not reported consistently in other studies (Devolder et al., 2024; Gilles et al., 2016; Guest et al., 2017; Morse & Vander Werff, 2023; Sendesen & Turkyilmaz, 2025; Shim et al., 2017). Mixed results were also reported with envelope following responses (EFRs), an auditory evoked potential considered an indicator of cochlear synaptopathy (Garrett et al., 2025; Keshishzadeh et al., 2020; Shaheen, Valero, & Liberman, 2015; Vasilkov et al., 2021). Bramhall et al. (2023) reported a link between EFR and tinnitus, but this has not been confirmed by others (Devolder et al., 2024; Guest et al., 2017).

Furthermore, efferent neural activity has been increasingly studied in the context of hidden hearing loss and tinnitus, and can be assessed via auditory reflexes. The middle ear muscle reflex (MEMR) stiffens the stapedius muscle in response to loud sounds and is, along with the EFR, considered a potential biomarker of cochlear synaptopathy (Mepani et al., 2019; Valero et al., 2018; Wojtczak, Beim, & Oxenham, 2017). Tinnitus-related increments in MEMR thresholds and reduced amplitudes were reported by several studies (Bramhall et al., 2023; da Cruz Fernandes et al., 2013; Vasilkov et al., 2023; Wojtczak, Beim, & Oxenham, 2017), while others reported no such differences (Fernandes et al., 2014; Guest, Munro, & Plack, 2019). Secondly, the medial olivocochlear reflex (MOCR) inhibits outer hair cell (OHC) motility via efferent pathways from the superior olivary complex, causing OAE suppression. Tinnitus-related findings are mixed, with some showing reduced suppression (Degeest et al., 2014), some stronger suppression (Vasilkov et al., 2023), and others observing no differences (Cheng et al., 2020; Tayade & Tucker, 2022).

The relationship between tinnitus and hidden sensorineural hearing loss remains unclear across several biomarkers. One contributing factor may be age, which is closely associated with hidden sensorineural hearing loss. Inconsistencies in age matching between test groups can confound tinnitus-related interpretations. Additional variability may arise from interactions between sensorineural mechanisms along the auditory pathway, particularly between afferent and efferent circuits. This study aims to clarify how auditory sensorineural encoding relates to tinnitus, by disentangling potential confounding factors and measurement interactions. Specifically, we: (1) Study tinnitus-specific differences in sensorineural auditory encoding and speech perception within age-and hearing-status-matched groups, evaluating both envelope and temporal fine structure processing using high-and low-frequency sound stimuli. (2) Explore tinnitus-specific differences in efferent neural auditory pathways by assessing auditory reflexes. (3) Identify tinnitus-related mechanisms by examining relationships between objective markers of hidden sensorineural hearing loss, behavioral speech perception, and subjective tinnitus-related parameters.

## Materials and methods

### 1. Subjects and tinnitus characteristics

A total of 113 participants were included. To enable a clear dissociation between tinnitus-, age-, and hearing-status-related effects, participants were allocated to six groups: young normal-hearing adults without tinnitus (yNH_CO) or with tinnitus (yNH_T), middle-aged/older normal-hearing adults without tinnitus (oNH_CO) or with tinnitus (oNH_T), and middle-aged/older adults with hearing loss without tinnitus (oHI_CO) or with tinnitus (oHI_T).

The young cohort comprised individuals aged 18 to 35 years, and the older middle-aged those aged 45 to 65 years. (Near-)normal hearing was defined by audiometric thresholds, with slightly looser criteria for high frequencies in older groups to accommodate minimal age-related threshold shifts: for younger groups ≤ 20 dB HL from 125 to 8000 Hz and for older groups ≤ 20 dB HL from 125 to 1000 Hz and ≤ 30 dB HL from 2000 to 8000 Hz (Figure 1). This minor audiometric difference can influence age-related effects, but will not affect tinnitus-related comparisons, as criteria are the same for control and tinnitus groups within each hearing-status category (yNH, oNH, oHI). Within each hearing-status category, tinnitus and control groups were matched for sex and age (±2 yrs; descriptive data in Table 1).

**Figure 1:**
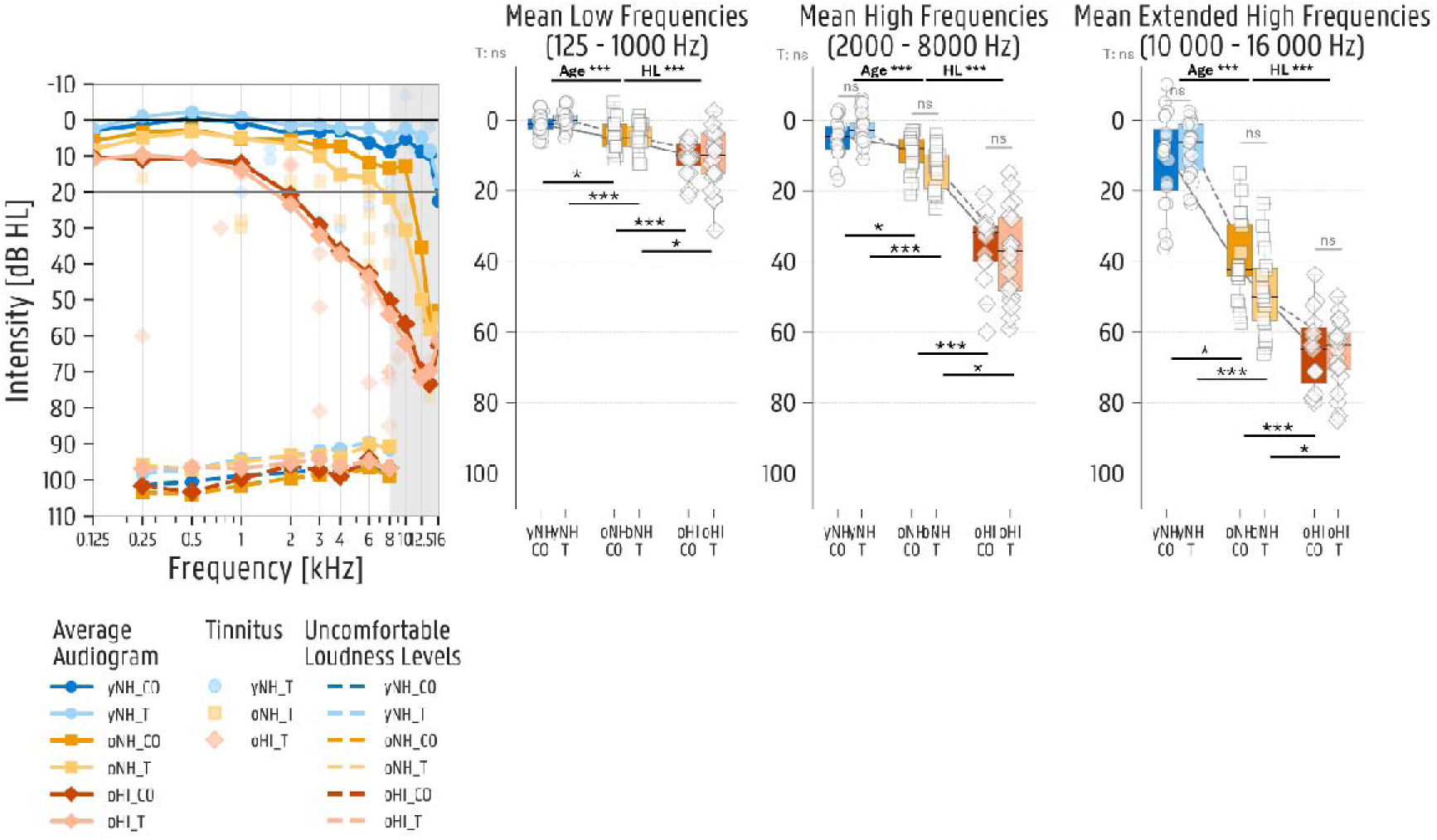
Average tonal audiometric thresholds, tinnitus loudness and pitch, and average uncomfortable loudness levels for the six test groups; yNH_CO = young normal hearing control (N=20), yNH_T = young normal hearing with tinnitus (N=20), oNH_CO = older normal hearing control (N=19), oNH_T = older normal hearing with tinnitus (N=19), oHI_CO = older hearing impaired control (N=15), oHI_T = older hearing impaired with tinnitus (N=20).

**Table 1:**
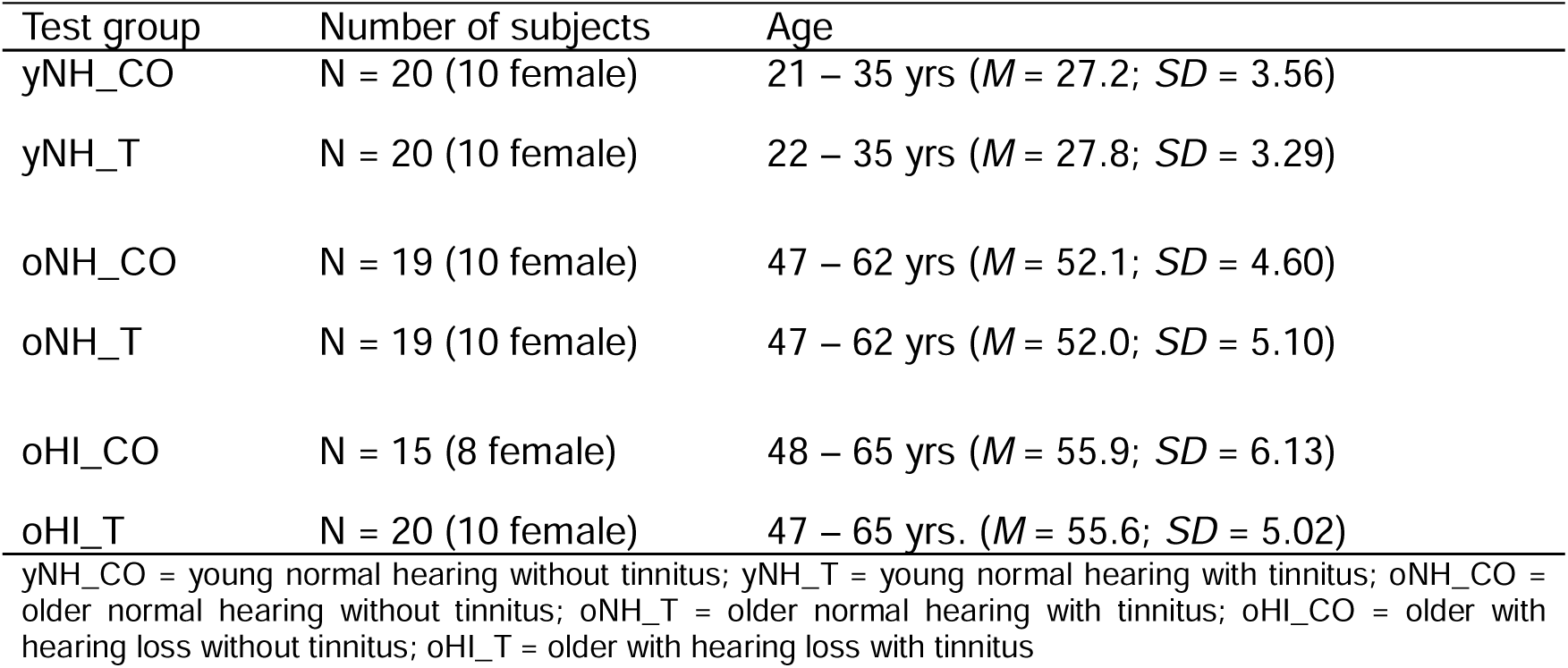
Descriptive characteristics of the six test groups.

| Test group | Number of subjects | Age |
|---|---|---|
| yNH_CO | N = 20 (10 female) | 21 – 35 yrs (*M* = 27.2; *SD* = 3.56) |
| yNH_T | N = 20 (10 female) | 22 – 35 yrs (*M* = 27.8; *SD* = 3.29) |
| oNH_CO | N = 19 (10 female) | 47 – 62 yrs (*M* = 52.1; *SD* = 4.60) |
| oNH_T | N = 19 (10 female) | 47 – 62 yrs (*M* = 52.0; *SD* = 5.10) |
| oHI_CO | N = 15 (8 female) | 48 – 65 yrs (*M* = 55.9; *SD* = 6.13) |
| oHI_T | N = 20 (10 female) | 47 – 65 yrs. (*M* = 55.6; *SD* = 5.02) |

yNH\_CO = young normal hearing without tinnitus; yNH\_T = young normal hearing with tinnitus; oNH\_CO = older normal hearing without tinnitus; oNH\_T = older normal hearing with tinnitus; oHI\_CO = older with hearing loss without tinnitus; oHI\_T = older with hearing loss with tinnitus

General inclusion criteria required participants to be free from major health conditions and to have Flemish Dutch as their native language. Exclusion criteria included any known middle-ear pathology or middle-ear surgery (except ventilation tubes), a history or current use of ototoxic medication, pregnancy, and the use of hearing aids or implants. Participants in the hearing loss groups presented exclusively with sensorineural, sloping audiometric configurations with an air–bone gap < 15 dB.

Tinnitus-related inclusion criteria were continuous, subjective, non-pulsatile tinnitus of chronic duration (> 1 year), with an etiology of noise-induced, age-related or idiopathic origin. Tinnitus-related exclusion criteria included middle-ear–related causes, head trauma, medication-induced tinnitus, and a history of tinnitus treatment involving neuromodulation or hearing aids. Participants reporting hyperacusis were excluded. Hyperacusis was evaluated based on the question “Are you oversensitive to sound? (= sounds that sound normal for others, are often uncomfortable or painful for me)”, or a score of 28 or above on the hyperacusis questionnaire (HQ). The HQ consists of 14 questions that can be answered with ‘not true’, ‘sometimes true’, ‘often true’, ‘always true’ (Khalfa et al., 2002; Meeus et al., 2010).

Besides the HQ, all participants completed questionnaires before the start of the experiment, covering the following topics: (i) general sociodemographic questions, (ii) subjective hearing difficulties, (iii) tinnitus and (iv) noise exposure and use of hearing protection. Participants with tinnitus completed the Dutch version of the following questionnaires assessing tinnitus characteristics and distress: the Tinnitus Sample Case History Questionnaire (TSCHQ) (Langguth et al., 2007), the Tinnitus Functional Index (TFI) (Meikle et al., 2012; Rabau, Wouters, & Van de Heyning, 2014), and the Tinnitus Handicap Inventory (THI) (Newman, Sandridge, & Bolek, 2008).

Tinnitus characteristics and questionnaire scores are summarized in Table 2. Tinnitus pitch and loudness were assessed using a standard tinnitus matching procedure, conducted with the same setup as for audiometry (Interacoustics Equinox audiometer with RadioEar DD450 headphones in a double-walled booth). Participants first chose the sound that best resembled their tinnitus, a pure tone or a narrowband noise. Frequency pairs were then presented until the pitch most closely matching the tinnitus was identified. Loudness at the matched pitch was assessed beginning at 10 dB SL, with intensity adjusted in 1 dB increments based on participant feedback. Minimal masking levels were assessed using the same speech-shaped noise as in the speech in noise (SPIN) test, delivered through a Fireface UCX soundcard (RME) and HDA-300 Sennheiser headphones via MATLAB R2023b. The noise intensity increased in 5 dB steps until the participant reported that their tinnitus was masked.

**Table 2:**
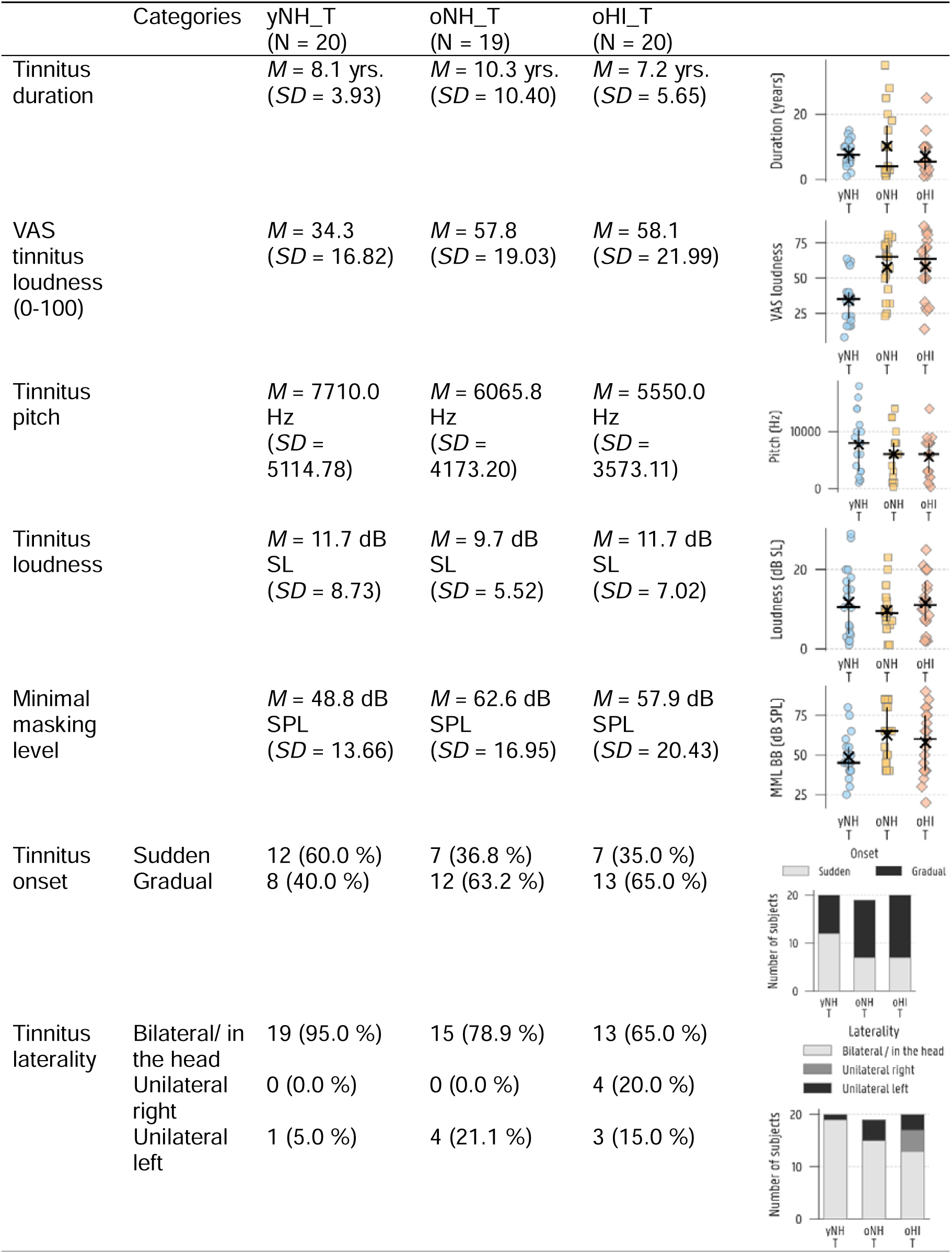
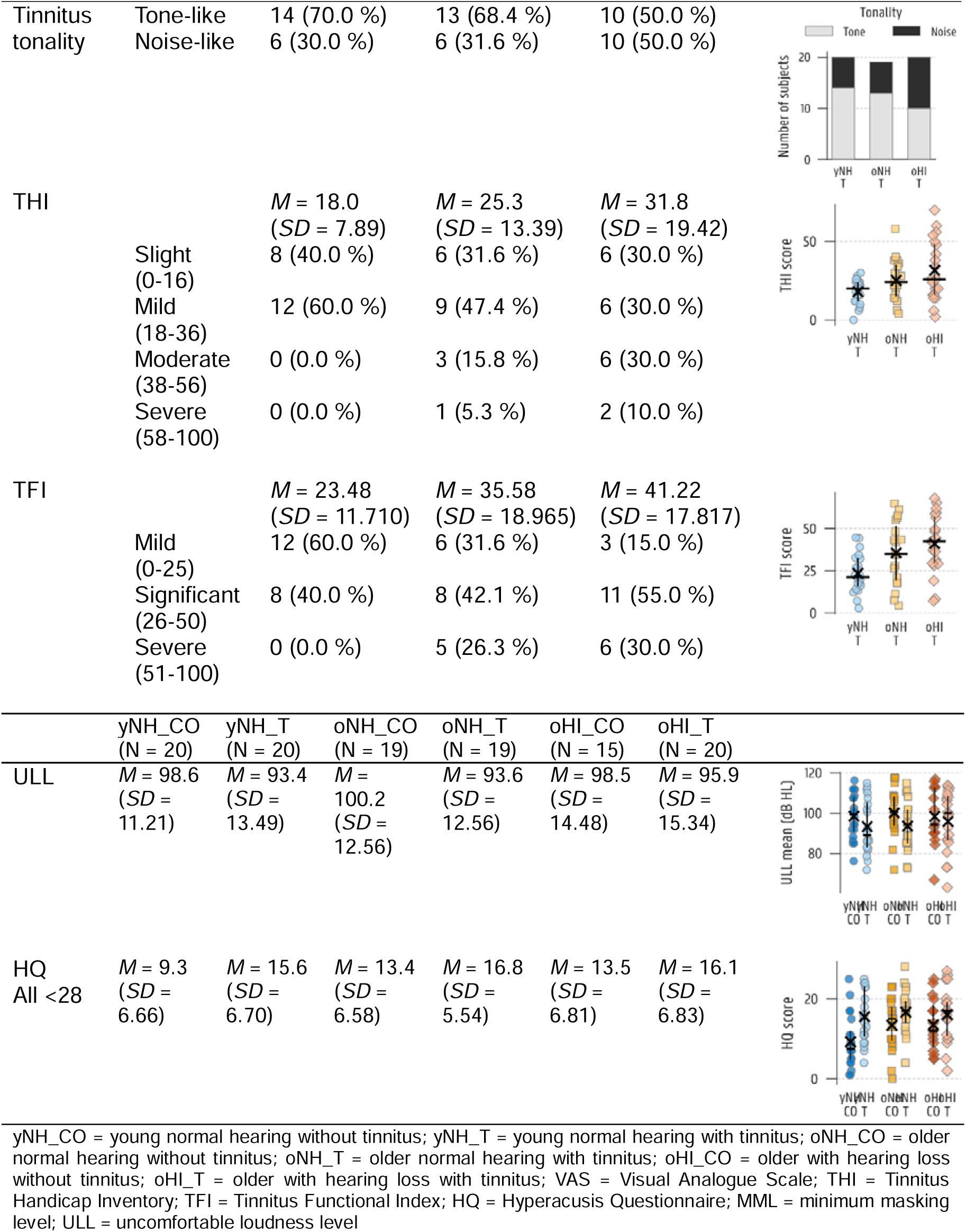
Descriptive tinnitus characteristics and questionnaire results of the three test groups with tinnitus and hyperacusis-related parameters for all six test groups.

|  | Categories | yNH_T<br>(N = 20) | oNH_T<br>(N = 19) | oHI_T<br>(N = 20) |  |  |
|---|---|---|---|---|---|---|
| Tinnitus duration |  | *M* = 8.1 yrs.<br>(*SD* = 3.93) | *M* = 10.3 yrs.<br>(*SD* = 10.40) | *M* = 7.2 yrs.<br>(*SD* = 5.65) |  |  |
| VAS tinnitus loudness (0-100) |  | *M* = 34.3<br>(*SD* = 16.82) | *M* = 57.8<br>(*SD* = 19.03) | *M* = 58.1<br>(*SD* = 21.99) |  |  |
| Tinnitus pitch |  | *M* = 7710.0 Hz<br>(*SD* = 5114.78) | *M* = 6065.8 Hz<br>(*SD* = 4173.20) | *M* = 5550.0 Hz<br>(*SD* = 3573.11) |  |  |
| Tinnitus loudness |  | *M* = 11.7 dB SL<br>(*SD* = 8.73) | *M* = 9.7 dB SL<br>(*SD* = 5.52) | *M* = 11.7 dB SL<br>(*SD* = 7.02) |  |  |
| Minimal masking level |  | *M* = 48.8 dB SPL<br>(*SD* = 13.66) | *M* = 62.6 dB SPL<br>(*SD* = 16.95) | *M* = 57.9 dB SPL<br>(*SD* = 20.43) |  |  |
| Tinnitus onset | Sudden | 12 (60.0 %) | 7 (36.8 %) | 7 (35.0 %) |  |  |
|  | Gradual | 8 (40.0 %) | 12 (63.2 %) | 13 (65.0 %) |  |  |
| Tinnitus laterality | Bilateral/ in the head | 19 (95.0 %) | 15 (78.9 %) | 13 (65.0 %) |  |  |
|  | Unilateral right | 0 (0.0 %) | 0 (0.0 %) | 4 (20.0 %) |  |  |
|  | Unilateral left | 1 (5.0 %) | 4 (21.1 %) | 3 (15.0 %) |  |  |
| Tinnitus tonality | Tone-like | 14 (70.0 %) | 13 (68.4 %) | 10 (50.0 %) |  |  |
|  | Noise-like | 6 (30.0 %) | 6 (31.6 %) | 10 (50.0 %) |  |  |
| THI |  | *M* = 18.0<br>(*SD* = 7.89) | *M* = 25.3<br>(*SD* = 13.39) | *M* = 31.8<br>(*SD* = 19.42) |  |  |
|  | Slight (0-16) | 8 (40.0 %) | 6 (31.6 %) | 6 (30.0 %) |  |  |
|  | Mild (18-36) | 12 (60.0 %) | 9 (47.4 %) | 6 (30.0 %) |  |  |
|  | Moderate (38-56) | 0 (0.0 %) | 3 (15.8 %) | 6 (30.0 %) |  |  |
|  | Severe (58-100) | 0 (0.0 %) | 1 (5.3 %) | 2 (10.0 %) |  |  |
| TFI |  | *M* = 23.48<br>(*SD* = 11.710) | *M* = 35.58<br>(*SD* = 18.965) | *M* = 41.22<br>(*SD* = 17.817) |  |  |
|  | Mild (0-25) | 12 (60.0 %) | 6 (31.6 %) | 3 (15.0 %) |  |  |
|  | Significant (26-50) | 8 (40.0 %) | 8 (42.1 %) | 11 (55.0 %) |  |  |
|  | Severe (51-100) | 0 (0.0 %) | 5 (26.3 %) | 6 (30.0 %) |  |  |
|  | yNH_CO<br>(N = 20) | yNH_T<br>(N = 20) | oNH_CO<br>(N = 19) | oNH_T<br>(N = 19) | oHI_CO<br>(N = 15) | oHI_T<br>(N = 20) |
| ULL | *M* = 98.6<br>(*SD* = 11.21) | *M* = 93.4<br>(*SD* = 13.49) | *M* = 100.2<br>(*SD* = 12.56) | *M* = 93.6<br>(*SD* = 12.56) | *M* = 98.5<br>(*SD* = 14.48) | *M* = 95.9<br>(*SD* = 15.34) |
| HQ<br>All <28 | *M* = 9.3<br>(*SD* = 6.66) | *M* = 15.6<br>(*SD* = 6.70) | *M* = 13.4<br>(*SD* = 6.58) | *M* = 16.8<br>(*SD* = 5.54) | *M* = 13.5<br>(*SD* = 6.81) | *M* = 16.1<br>(*SD* = 6.83) |

yNH_CO = young normal hearing without tinnitus; yNH_T = young normal hearing with tinnitus; oNH_CO = older normal hearing without tinnitus; oNH_T = older normal hearing with tinnitus; oHI_CO = older with hearing loss without tinnitus; oHI_T = older with hearing loss with tinnitus; VAS = Visual Analogue Scale; THI = Tinnitus Handicap Inventory; TFI = Tinnitus Functional Index; HQ = Hyperacusis Questionnaire; MML = minimum masking level; ULL = uncomfortable loudness level

The study was approved by the ethical committee at Ghent University (ONZ-2022-0352 E01). All participants provided written informed consent prior to enrollment.

### 2. Hearing status and uncomfortable loudness levels

Tympanometry was performed using the Sentiero Tymp Diagnostic (Path Medical) with a 226-Hz probe tone at 85 dB SPL. Pure-tone audiometry was conducted in a double-walled booth using an Interacoustics Equinox audiometer and RadioEar DD450 headphones, following the modified Hughson-Westlake method. Air-conduction thresholds were measured at 0.125, 0.250, 0.500, 1, 2, 3, 4, 6, and 8 kHz, and at extended high frequencies of 10, 12.5, 14, and 16 kHz. When thresholds between 0.5 and 4 kHz exceeded 20 dB HL, bone-conduction thresholds were obtained. Participants with an air–bone gap ≥ 15 dB were excluded. For all remaining procedures, the better-hearing ear was tested, except in cases of unilateral or significantly asymmetric tinnitus, where the ear in which tinnitus was most prominent was selected.

For the analyses, audiometric thresholds of the tested ear were summarised in three parameters: Audio_LF (mean of 0.125, 0.25, 0.5, and 1 kHz), Audio_HF (mean of 2, 3, 4, 6, and 8 kHz), and Audio_EHF (mean of 10, 12.5, 14, and 16 kHz). For participants with tinnitus, an additional parameter, Audio_Tin, was defined as the threshold at the tinnitus pitch.

Uncomfortable loudness levels (ULLs, Figure 1, Table 2) were measured using the same audiometric setup. Pure-tone stimuli were presented at 250, 500, 1000, 2000, 3000, 4000, 6000, and 8000 Hz, beginning at 60 dB HL and increasing in 5 dB steps (BSA, 2011). Participants indicated when the sound became uncomfortably loud by pressing a response button.

### 3. Measures of sensorineural encoding

#### 3.1. Distortion product otoacoustic emissions (DPOAE)

OHC integrity was assessed using DPOAEs with the Universal Smart Box (Intelligent Hearing Systems) and SmartDPOAE software. Stimuli were delivered through an Etymotic ER10D probe. Two primary tones were presented simultaneously at a frequency ratio f2/f1 = 1.22, with f2 ranging from 0.5 to 8 kHz at two frequencies per octave. Primary levels were L1 = 65 dB SPL and L2 = 55 dB SPL. Responses were recorded over 32 sweeps. Amplitudes falling below the noise floor were replaced with the noise floor value to prevent unreliable negative values from skewing the data. Mean amplitudes were calculated separately for low (0.5 – 1 kHz) and high (2 – 8 kHz) frequency ranges.

#### 3.2. Auditory brainstem response (ABR)

To examine neural sound transmission along the afferent auditory pathway, ABRs were collected at 70 and 90 dB peSPL using 80-µs clicks presented at a rate of 11 Hz, averaged across 3000 repetitions (Intelligent Hearing System with Universal Smart Box and SmartEP Continuous Acquisition Module). Electrodes were placed at Fz (positive), the earlobes (negative), and the nose (ground). Recordings were band-pass filtered between 100 and 1500 Hz, epoched, and baseline-corrected. Waves I, III, V, and VI were manually identified by trained audiologists to determine peak-to-trough amplitudes; corresponding to activity in the distal auditory nerve, superior olivary complex, inferior colliculus, and lateral lemniscus respectively. The wave I/V ratio (wave I amplitude divided by wave V amplitude) was analyzed as an exploratory complement to the separate wave I and wave V analyses.

#### 3.3. Frequency following response (FFR)

Using the same setup as for ABR, sensorineural frequency encoding was assessed with a 170-ms vowel /u/ (Gaudrain, Verhulst, & Başkent, 2025), with peaks at 116 Hz and its harmonics (highest at 232 Hz), similar to Dapper et al. (2025). Vowels were presented at 70 dB SPL at a rate of 4 Hz and averaged across 2500 sweeps. All FFR recordings were band-pass filtered (60–2000 Hz), baseline-corrected, and epoched, with epochs exceeding three times the median peak-to-peak amplitude rejected. Analysis followed the approach described in Ponsot et al. (2025). ENV and TFS components were derived from the sum and difference, respectively, of recordings to stimuli presented in alternating polarities (Aiken & Picton, 2008; Krishnan, 2002). Frequency spectra were noise-floor corrected via a bootstrap procedure, and the magnitudes of the first 10 harmonics were summed as outcome parameter.

#### 3.4. Envelope following response (EFR)

EFRs were obtained to assess temporal envelope encoding and potential cochlear synaptopathy, using the same recording setup as ABR and FFR. Stimuli consisted of a 4000 Hz carrier tone, amplitude-modulated with a 110 Hz rectangular waveform (RAM), presented in alternating polarity. Each stimulus was 490 ms in duration, presented at 70 dB SPL with a repetition rate of 2 Hz. A total of 800 sweeps were recorded per condition. Signals were band-pass filtered (30–1500 Hz), epoched, baseline-corrected, and analyzed using a bootstrapping approach to estimate the noise floor (Van Der Biest et al., 2023; Vasilkov et al., 2021). The EFR magnitude was calculated as the sum of the signal-to-noise ratio at the fundamental modulation frequency and the next three harmonics (110, 220, 330, 440 Hz).

#### 3.5. Middle ear muscle reflex (MEMR)

MEMR measurements were conducted in a sound-attenuating booth using an Etymotic ER-10X probe system connected to an RME Fireface UCX sound card and MATLAB R2023b software. Ipsilateral and contralateral MEMRs were elicited with broadband noise (0.5–8.5 kHz, 40–90 dB SPL in 5-dB steps) and measured using 94-µs clicks at 90 dB peSPL. Each trial consisted of seven probe clicks alternating with 120-ms noise bursts, repeated 16 times per ear with a 1.5-s intertrial interval (as in Bharadwaj et al. (2022)). Outlier trials exceeding two SDs from the mean were excluded. MEMR growth functions were baseline-corrected using responses at 40–50 dB SPL and fit with a three-parameter sigmoid function. Thresholds were defined as the elicitor level at which the fitted curve reached 0.02 dB above baseline. Poor fits (R² < 0.6) were excluded unless extremely shallow growth indicated an absent response. Thresholds exceeding 90 dB SPL or showing negligible growth were set to 95 dB SPL to indicate an elevated reflex.

#### 3.6. Medial olivocochlear reflex (MOCR)

MOCR recordings were obtained using the same setup as MEMR, but assessed changes in OAEs rather than ear canal pressure. Click-evoked OAEs (93.75 µs, 55 dB peSPL) were presented in a forward-masking paradigm using broadband noise elicitors (60 dB SPL, 478 ms) presented ipsilaterally, contralaterally, and bilaterally. Each trial comprised five sequential blocks (baseline, ipsilateral, contralateral, bilateral, baseline), with 400 trials collected per condition (as in Boothalingam et al. (2019)). Click evoked otoacoustic emission (CEOAE) responses (5–20 ms post-click) were band-pass filtered (0.6–6 kHz) and epochs exceeding two SDs of the mean RMS were rejected. Responses were split into two buffers to calculate CEOAE magnitude and noise; measurements with <80% reproducibility at baseline were excluded. MOCR-induced suppression (ΔCEOAE) was quantified as the reduction in CEOAE spectral amplitude in elicitor condition relative to baseline.

### 4. Speech-related outcome measurements

#### 4.1. Vowel discrimination task

An /o/-/u/ vowel pair with a fundamental frequency of 116 Hz was used, matching that of the FFR stimuli. The vowels differed only in the first formant (F1, <1.5 kHz) and were matched in duration and level to the /u/ vowel used in the FFR recordings (170 ms, 70 dB SPL) (further details in Gaudrain, Verhulst and Başkent (2025), Dapper et al. (2025) and Schirmer et al. (2024)). Stimuli were presented using a Fireface UCX soundcard (RME), HDA-300 Sennheiser headphones, and MATLAB R2023b. A three-interval, odd-one-out procedure was conducted, with two contrast widths (large and small) selected to minimize ceiling and floor effects. The task was performed in quiet and with speech-shaped noise presented ipsilaterally at 0 dB SNR, with nine repetitions per contrast. Each condition began with four training trials, and no feedback was provided during the main experiment.

#### 4.2. Matrix sentence test

Speech intelligibility was measured using the Flemish Matrix sentence test (Luts et al., 2014), presented through the same setup as the vowel discrimination task. The test consisted of closed-set, unpredictable five-word sentences. Speech-shaped noise was fixed at 70 dB SPL, with an onset 500 ms before and an offset 500 ms after each sentence. Speech level was adjusted using a one-up, one-down procedure to track the 50% speech reception threshold (SRT), starting at 50 dB SPL for speech in quiet (SPIQ) and 70 dB SPL for SPIN (0 dB SNR at onset). The initial step size was 5 dB, decreasing to less than 1 dB near threshold. Three SPIN conditions were tested; with broadband (BB), low-pass (LP, 1500 Hz cutoff), and high-pass (HP, 1650 Hz cutoff) filtering applied to both speech and noise using a 1,024th-order FIR filter. BB, LP, and HP SPIN conditions were each measured twice using two interleaved adaptive tracks, while SPIQ (BB, LP, HP) were administered once. A brief training block (BB SPIQ and BB SPIN) preceded testing.

### 5. Statistical analysis

All statistical analyses were conducted in Python 3.11.4. Group comparisons and correlation analyses were performed to distinguish tinnitus-, age-and hearing-loss-related effects, including their potential interactions.

Multiple ANOVAs (α = 0.05) were conducted for each dependent variable, with planned post-hoc comparisons. Effects were assessed in three separate models (α = 0.05) for each dependent variable:

• Tinnitus (2 x 3 ANOVA): tinnitus (yNH_T, oNH_T, oHI_T) vs. control (yNH_CO, oNH_CO, oHI_CO)

Planned post-hoc comparisons if the main effect is significant:

`o` yNH_T vs. yNH_CO
`o` oNH_T vs. oNH_CO
`o` oHI_T vs. oHI_CO

• Age (2 x 2 ANOVA): young (yNH_CO, yNH_T) vs. older (oNH_CO, oNH_T)

Planned post-hoc comparisons if the main effect or interaction is significant:

`o` yNH_CO vs. oNH_CO
`o` yNH_T vs. oNH_T

• Audiometric Hearing Loss (2 x 2 ANOVA): normal hearing (oNH_CO, oNH_T) vs. hearing loss (oHI_CO, oHI_T)

Planned post-hoc comparisons if the main effect or interaction is significant:

`o` oNH_CO vs. oHI_CO
`o` oNH_T vs. oHI_T

Normality of residuals and homogeneity of variances were assessed with Shapiro–Wilk and Levene’s tests, respectively. When residuals deviated from normality, a power transformation was applied to reduce skewness: Box–Cox transformation was used for strictly positive data, whereas Yeo–Johnson transformation was used when values included zero or negative observations. Assumptions were reassessed after each transformation. A transformation was retained if it satisfied both assumptions or restored residual normality while leaving heteroscedasticity that could be addressed using the procedures described below. If residuals were normally distributed but unequal variances persisted, ANOVA with HC3 heteroscedasticity-consistent standard errors was used, and planned pairwise comparisons were conducted using Welch’s t-tests. If residual non-normality persisted, aligned rank transform (ART) ANOVA was used, with Bonferroni-corrected Mann–Whitney U tests for planned pairwise comparisons.

Effect sizes are reported as partial eta-squared (η²⍰). Planned post-hoc comparisons were performed using Student’s t-tests (or Welch’s t-tests or Mann-Whitney U tests, as above), with Bonferroni correction applied for the number of comparisons (Tinnitus: 3; Age: 2; Hearing Loss: 2; reported p-values are corrected). Effect sizes for post-hoc tests were Cohen’s d (t-tests) or the rank-biserial correlation (Mann-Whitney). Main effect sizes and significance across all measures were summarized in a factor-by-variable heatmap. Means and standard deviations of the six groups are provided in Table S1 for all included variables.

To account for potential confounding effects of OHC damage, ANOVAs were repeated with variables residualized against DPOAE amplitudes, thereby isolating effects independent of peripheral cochlear amplification (ANCOVA). All dependent variables were residualized against mean low-frequency and high-frequency DPOAE amplitudes.

Correlations were restricted to normal-hearing participants (yNH_CO, yNH_T, oNH_CO, oNH_T) and to combined groups (NH_Control = yNH_CO + oNH_CO; NH_Tinnitus = yNH_T + oNH_T) to avoid confounds introduced by audiometric hearing loss. All variables were residualized against mean low-and high-frequency DPOAE amplitudes prior to correlation. For each pair of variables and each (sub)group, normality of both variables was assessed with the Shapiro–Wilk test (α = 0.01). Pearson correlations (r) were computed when both variables were normally distributed, and Spearman rank correlations (ρ) when at least one variable deviated from normality. Correlations were considered significant at α = 0.01, to strike a balance between the exploratory nature and the risk of inflated Type I error due to multiple comparisons.

## Results

Mean audiometric thresholds revealed no significant main effect of tinnitus (*p* >.05; Figure 1). Age-related effects were present at low frequencies (LFs; *F*(1,74) = 28.43, *p* <.001, η²⍰ = 0.28), high frequencies (HFs; *F*(1,74) = 40.92, *p* <.001, η²⍰ = 0.36) and extended high frequencies (EHFs; *F*(1,74) = 172.54, *p* <.001, η²⍰ = 0.70), with oNH groups exhibiting higher thresholds than yNH groups. Significant hearing-loss-related effects were observed across all three frequency bands: LF (*F*(1,69) = 23.36, *p* <.001, η²⍰ = 0.25), HF (*F*(1,69) = 134.62, *p* <.001, η²⍰ = 0.66) and EHF (*F*(1,69) = 68.64, *p* <.001, η²⍰ = 0.50). An interaction between age and tinnitus was observed for HFs (*F*(1,74) = 9.40, *p* =.003, η²⍰ = 0.11), where the oNH_T group visually showed non-significant worse thresholds compared to the oNH_CO group, while the reverse pattern was apparent in the younger groups (Figure 1). A significant age-by-tinnitus interaction was also found for EHFs (*F*(1,74) = 6.01, *p* =.017, η²⍰ = 0.08).

Hyperacusis was excluded based on HQ scores (below 28). Nevertheless, subjects with tinnitus had significantly higher HQ scores than control subjects (*F*(1,109) = 11.37, *p* = 0.001, η²⍰ = 0.09, post-hoc yNH_CO vs. yNH_T: *t*(38) =-2.96, *p_Bonf_* =.016, *d* =-0.94). Also mean ULL’s showed a weak significant difference between the tinnitus and control groups (*F*(1,109) = 4.09, *p* =.046, η²⍰ = 0.04).

### 1. Sensorineural encoding

#### 1.1. Distortion product otoacoustic emissions (DPOAE)

Mean DPOAE amplitudes were significantly reduced as a function of both age and hearing loss (HL) in the LF range (Age: *F*(1,72) = 12.69, *p* <.001, η²⍰ = 0.15; HL: *F*(1,68) = 8.35, *p* =.005, η²⍰ = 0.11; post-hoc yNH_CO vs oNH_CO: *t*(37) = 2.54, *p*_Bonf_ =.031, *d* = 0.81; post-hoc yNH_T vs oNH_T: *t*(35) = 2.51, *p*_Bonf_ =.034, *d* = 0.83; post-hoc oNH_T vs oHI_T: *t*(36) = 2.75, *p*_Bonf_ =.018, *d* = 0.90) and the HF range (Age: *F*(1,72) = 15.67, *p* <.001, η²⍰ = 0.18; HL: *F*(1,68) = 43.02, *p* <.001, η²⍰ = 0.39; post-hoc yNH_CO vs oNH_CO: *t*(37) = 2.92, *p*_Bonf_ =.012, *d* = 0.94; post-hoc yNH_T vs oNH_T: *t*(35) = 2.68, *p*_Bonf_ =.022, *d* = 0.88; post-hoc oNH_CO vs oHI_CO: *t*(32) = 5.09, *p*_Bonf_ <.001, *d* = 1.76; post-hoc oNH_T vs oHI_T: *t*(36) = 4.26, *p*_Bonf_ <.001, *d* = 1.38; Table 3; Figure S1). No significant tinnitus-related main effects or interactions emerged (*p* >.05), indicating no measurable differences in OHC function between control and tinnitus groups. These findings underscore the importance of distinguishing test groups by age and hearing status in subsequent analyses.

**Table 3:** Heatmap showing partial eta-squared effect sizes (η²p) for the influence of tinnitus status, age, hearing loss, and their interactions on each dependent variable. Each row represents a measurement outcome, and each column represents a main predictor or interaction term. *p <.05; **p <.01; ***p <.001. T = tinnitus, HL = Hearing Loss, DPOAE = Distortion Product Otoacoustic Emissions, ABR = Auditory Brainstem Response, amp = amplitude, lat = latency, EFR = Envelope Following Response, FFR = Frequency Following Response, ENV = Envelope, TFS = Temporal Fine Structure.

|  | Tinnitus | Age | Hearing Loss | T × Age | T × HL |
|---|---|---|---|---|---|
| DPOAE low frequencies |  | *** | ** |  |  |
| DPOAE high frequencies |  | *** | *** |  |  |
| ABR wave I amp 70 dB peSPL |  |  |  |  |  |
| ABR wave I amp 90 dB peSPL |  | ** |  |  |  |
| ABR wave V amp 70 dB peSPL |  | *** |  |  |  |
| ABR wave V amp 90 dB peSPL |  | ** |  | * |  |
| ABR wave I/V 70 dB peSPL |  |  |  |  |  |
| ABR wave I/V 90 dB peSPL | * |  |  | ** | * |
| ABR wave I lat 70 dB peSPL |  |  |  |  |  |
| ABR wave I lat 90 dB peSPL |  |  |  |  |  |
| ABR wave V lat 70 dB peSPL |  | *** | * |  |  |
| ABR wave V lat 90 dB peSPL |  | *** |  |  |  |
| /U/-FFR ENV 70 dB SPL |  | *** |  |  |  |
| /U/-FFR TFS 70 dB SPL |  | *** |  |  |  |
| EFR 4 kHz 70 dB SPL |  | *** | *** | * |  |

#### 1.2. Auditory brainstem response (ABR)

At 90 dB peSPL (Figure 2), significant age-related reductions were found in ABR wave I (*F*(1,72) = 10.48, *p* =.002, η²⍰ = 0.13; Table 3) and wave V amplitudes (*F*(1,72) = 9.82, *p* =.002, η²⍰ = 0.12). Post-hoc comparisons revealed a significant wave I difference between yNH_CO and oNH_CO (*t*(35) = 3.11, *p*_Bonf_ =.007, *d* = 1.02) and a significant wave V difference between yNH_T and oNH_T (*t*(37) = 4.66, *p_Bonf_* <.001, *d* = 1.49). For wave V, similar to HF and EHF audiometric thresholds, a significant Age × Tinnitus interaction was present (*F*(1,72) = 6.60, *p* =.012, η²⍰ = 0.08; Table 3), driven by a trend toward higher amplitudes in the yNH_T group relative to yNH_CO, and lower amplitudes in the oNH_T groups relative to oNH_CO. Although tinnitus had no significant main effect on ABR wave amplitudes (*p* >.05), it significantly affected the ABR wave I/V ratio (*F*(1,107) = 5.02, *p* =.027, η²⍰ = 0.05), with lower ratios in the tinnitus groups, consistent with, but not conclusive for, increased central gain. Post-hoc analysis revealed a significant tinnitus-related effect in the yNH groups (*U* = 294.0, *p*_Bonf_ =.011, *r*_rb_ = 0.55). Significant Tinnitus x Age (*F*(1,72) = 10.62, *p* =.002, η²⍰ = 0.13) and Tinnitus x Hearing Loss (*F*(1,66) = 4.30, *p* =.042, η²⍰ = 0.06) interactions were also observed for the wave I/V ratio (post-hoc yNH_CO vs oNH_CO: *U* = 255.0, *p*_Bonf_ =.022, *r*_rb_ = 0.49). Regarding ABR latencies, a significant age-related effect was observed for wave V (*F*(1,72) = 15.51, *p* <.001, η²⍰ = 0.18; post-hoc yNH_T vs oNH_T: *U* = 81.5, *p*_Bonf_ =.005, *r*_rb_ =-0.57), with no significant interactions.

**Figure 2:**
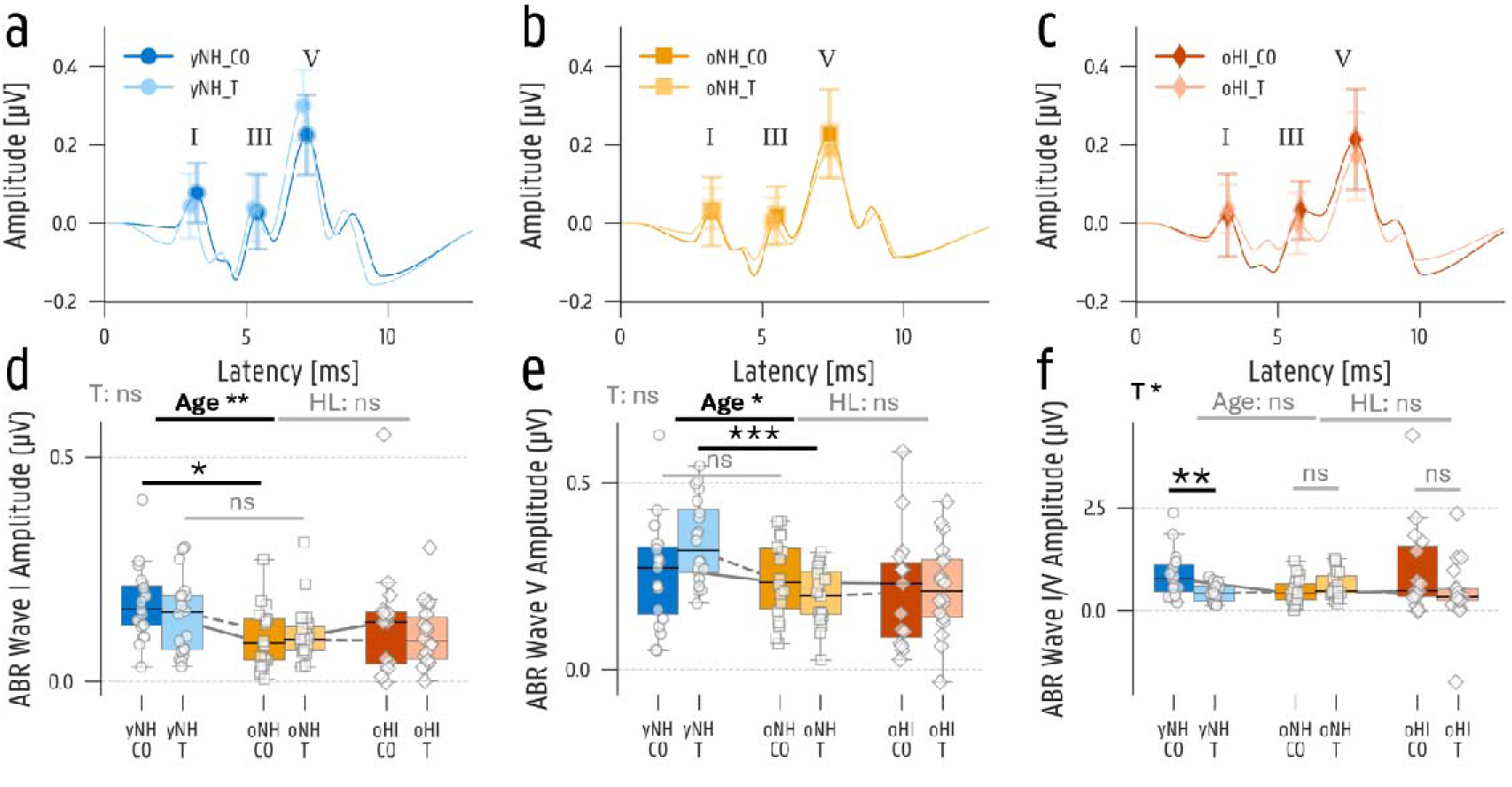
Auditory brainstem responses (ABR) with a 90 dB peSPL click. (a, b, c) Mean ABR-latency and-amplitude values for each peak-picked peak and trough (P1, N1, N2, P3, N3, P5, N5, P6, N6) for young normal hearing, older normal hearing and hearing impaired groups. Means are presented for each test group, and connected with light, curved trend lines to improve clarity. (d, e, f) Boxplot of the wave I, wave V and wave I/V ratio peak-to-trough amplitudes for each test group. yNH_CO = young normal hearing control, yNH_T = young normal hearing with tinnitus, oNH_CO = older normal hearing control, oNH_T = older normal hearing with tinnitus, oHI_CO = older hearing impaired control, oHI_T = older hearing impaired with tinnitus. *p <.05; **p <.01; ***p <.001, ns = not significant.

At 70 dB peSPL, significant age-related effects were limited to wave V amplitude (*F*(1,72) = 12.00, *p* <.001, η²⍰ = 0.14; post-hoc yNH_T vs oNH_T: *t*(37) = 3.83, *p_Bonf_* =.001, *d* = 1.23; Table 3) and latency (*F*(1,72) = 22.25, *p* <.001, η²⍰ = 0.24; post-hoc yNH_CO vs oNH_CO: *U* = 75.5, *p*_Bonf_ =.008, *r*_rb_ =-0.56; post-hoc yNH_T vs oNH_T: *U* = 86.0, *p*_Bonf_ =.007, *r*_rb_ =-0.55). One hearing-loss related effect was found in wave V latency (F(1,67) = 4.86, *p* =.031, η²⍰ = 0.07). No significant effects were observed for wave I amplitude or the wave I/V ratio (*p* >.05).

#### 1.3. Frequency following response (FFR)

FFR magnitudes for the /u/ stimulus revealed significant age-related main effects for both the envelope (*F*(1,72) = 22.32, *p* <.001, η²⍰ = 0.24; post-hoc yNH_T vs oNH_T: *t*(37) = 5.04, *p_Bonf_* <.001, *d* = 1.61; Table 3; Figure S2) and temporal fine structure components (*F*(1,72) = 41.40, *p* <.001, η²⍰ = 0.37; post-hoc yNH_CO vs oNH_CO: *t*(35) = 4.07, *p*_Bonf_ <.001, *d* = 1.34; yNH_T vs oNH_T: *t*(37) = 5.02, *p*_Bonf_ <.001, *d* = 1.61). No significant tinnitus-or hearing-loss-related effects were observed for either the ENV or TFS components (*p* >.05).

#### 1.4. Envelope following response (EFR)

Consistent with the proposed link with cochlear synaptopathy, a significant age-related reduction in EFR magnitude was observed (*F*(1,72) = 24.64, *p* <.001, η²⍰ = 0.25; Table 3). Hearing loss likewise exerted a significant effect on the EFR (*F*(1,68) = 16.90, *p* <.001, η²⍰ = 0.20), whereas no tinnitus-related main effects were found (*p* >.05). Post-hoc analyses are marked in Figure 3b, with significant differences between; yNH_T and oNH_T (*t*(37) = 5.31, *p_Bonf_* <.001, d = 1.70); oNH_CO and oHI_CO (*t*(31) = 3.34, *p_Bonf_* =.004, d = 1.17); oNH_T and oHI_T (*t*(37) = 2.71, *p_Bonf_* =.020, *d* = 0.87). A significant Tinnitus x Age interaction was found (*F*(1,72) = 6.61, *p* =.012, η²⍰ = 0.08), driven by a trend toward enhanced EFR magnitudes in the yNH_T group and a trend toward reduced magnitude in the oNH_T group (Figure 3).

**Figure 3:**
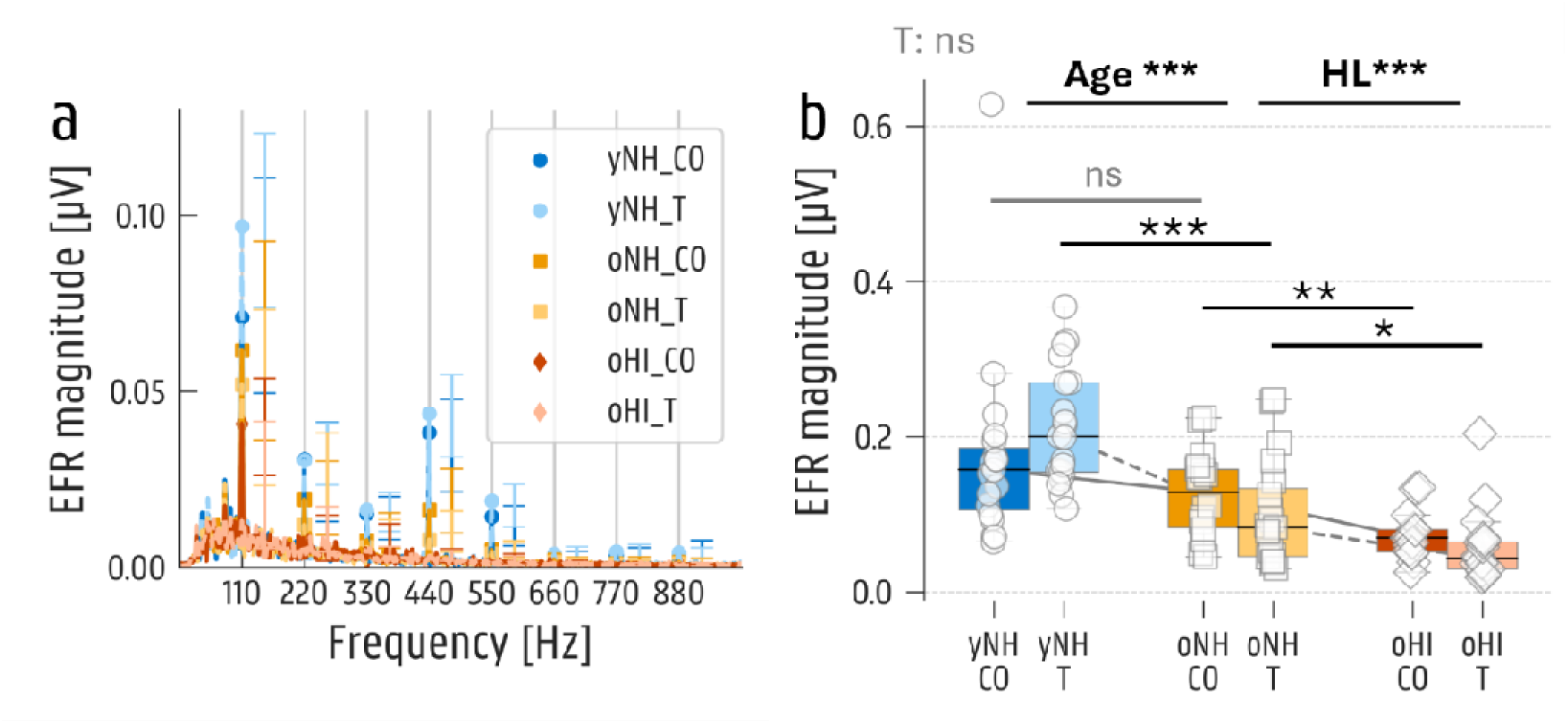
Envelope following response (EFR) with a 70 dB SPL rectangular amplitude modulated (110 Hz) tone of 4000 Hz. (a) Mean EFR response per test group with indication of the 110 Hz harmonic peaks. (b) Boxplot of the sum of the harmonic magnitudes for each test group. yNH_CO = young normal hearing control, yNH_T = young normal hearing with tinnitus, oNH_CO = older normal hearing control, oNH_T = older normal hearing with tinnitus, oHI_CO = older hearing impaired control, oHI_T = older hearing impaired with tinnitus. *p <.05; **p <.01; ***p <.001, ns = not significant.

#### 1.5. Middle ear muscle reflex (MEMR)

To examine potential differences in efferent pathways or cochlear synaptopathy, MEMR thresholds and amplitudes were analyzed (Table 4; Figure 4a). Significant age-related effects were observed for MEMR threshold (*F*(1,65) = 6.20, *p* =.015, η²⍰ = 0.09) and amplitude at 90 dB (*F*(1,74) = 9.34, *p* =.003, η²⍰ = 0.11). No other significant hearing-loss-or tinnitus-related differences were observed in this dataset.

**Figure 4:**
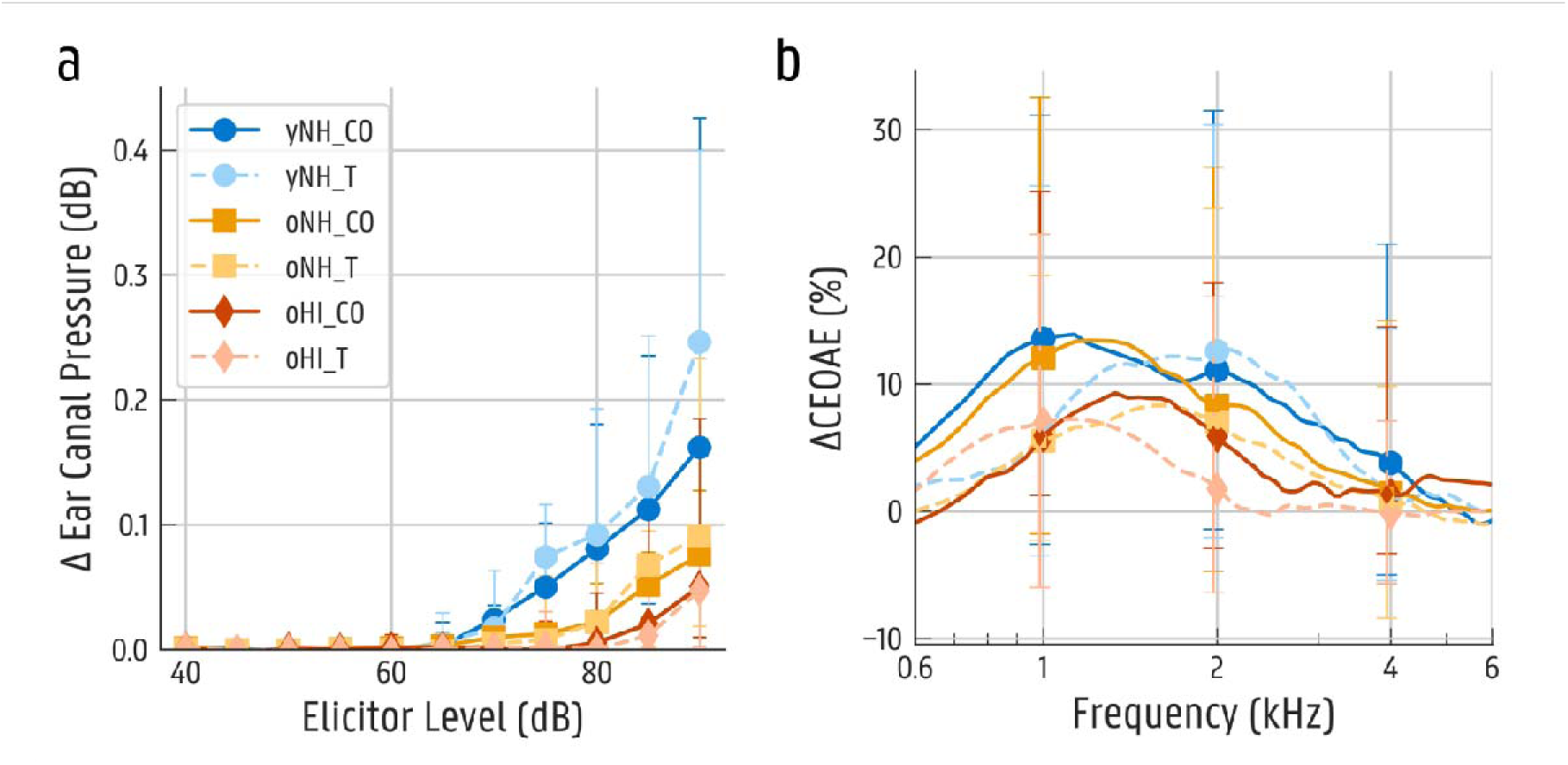
(a) Median ipsilateral middle ear muscle reflex (MEMR) growth function in ear canal pressure change and (b) ipsilateral medial olivocochlear reflex (MOCR) outcome in click evoked otoacoustic emission (CEOAE) change. yNH_CO = young normal hearing control, yNH_T = young normal hearing with tinnitus, oNH_CO = older normal hearing control, oNH_T = older normal hearing with tinnitus, oHI_CO = older hearing impaired control, oHI_T = older hearing impaired with tinnitus, CO = control groups, T = tinnitus groups.

**Table 4:** Heatmap showing partial eta-squared effect sizes (η²p) for the influence of tinnitus status, age, hearing loss, and their interactions on each dependent variable. Each row represents a measurement outcome, and each column represents a main predictor or interaction term. *p <.05; **p <.01; ***p <.001. T = tinnitus, HL = Hearing Loss, MEMR = Middle Ear Muscle Reflex, MOCR = Medial Olivocochlear Reflex, CEOAE = Click Evoked Otoacoustic Emissions.

|  | Tinnitus | Age | Hearing Loss | T × Age | T × HL |
|---|---|---|---|---|---|
| MEMR ipsi threshold |  | * |  |  |  |
| MEMR ipsi amplitude at 90 dB sPL |  | ** |  |  |  |
| MOCR ipsi ΔCEOAE |  |  |  |  |  |
| LF MOCR ipsi ΔCEOAE | * |  |  |  |  |
| HF MOCR ipsi ΔCEOAE |  | * |  |  |  |

### 1.6. Medial olivocochlear reflex (MOCR)

Analysis of MOCR strength revealed no significant main effects of age, hearing loss, tinnitus, nor their interactions (*p* >.05; Table 4; Figure 4b). When analyses were restricted to LF datapoints, a significant tinnitus-related effect emerged (*F*(1,47) = 5.01, *p* =.030, η²⍰ = 0.10), with stronger MOCR activity in the control group. Age-and hearing-loss-related comparisons were not significant (*p* >.05). In the HF range, MOCR revealed significant age-related reductions (*F*(1,47) = 4.63, *p* =.037, η²⍰ = 0.09), with no significant tinnitus-or hearing-loss-related effects.

### 2. Speech-related outcome

#### 2.1. Vowel discrimination task

A significant tinnitus-related effect emerged in the quiet condition (*F*(1,108) = 4.74, *p* =.032, η²⍰ = 0.04), with tinnitus groups outperforming their matched control groups (Table 5; Figure 5a). No other main effects, interactions, or effects in the noise condition were significant (*p* >.05; Figure 5b).

**Figure 5:**
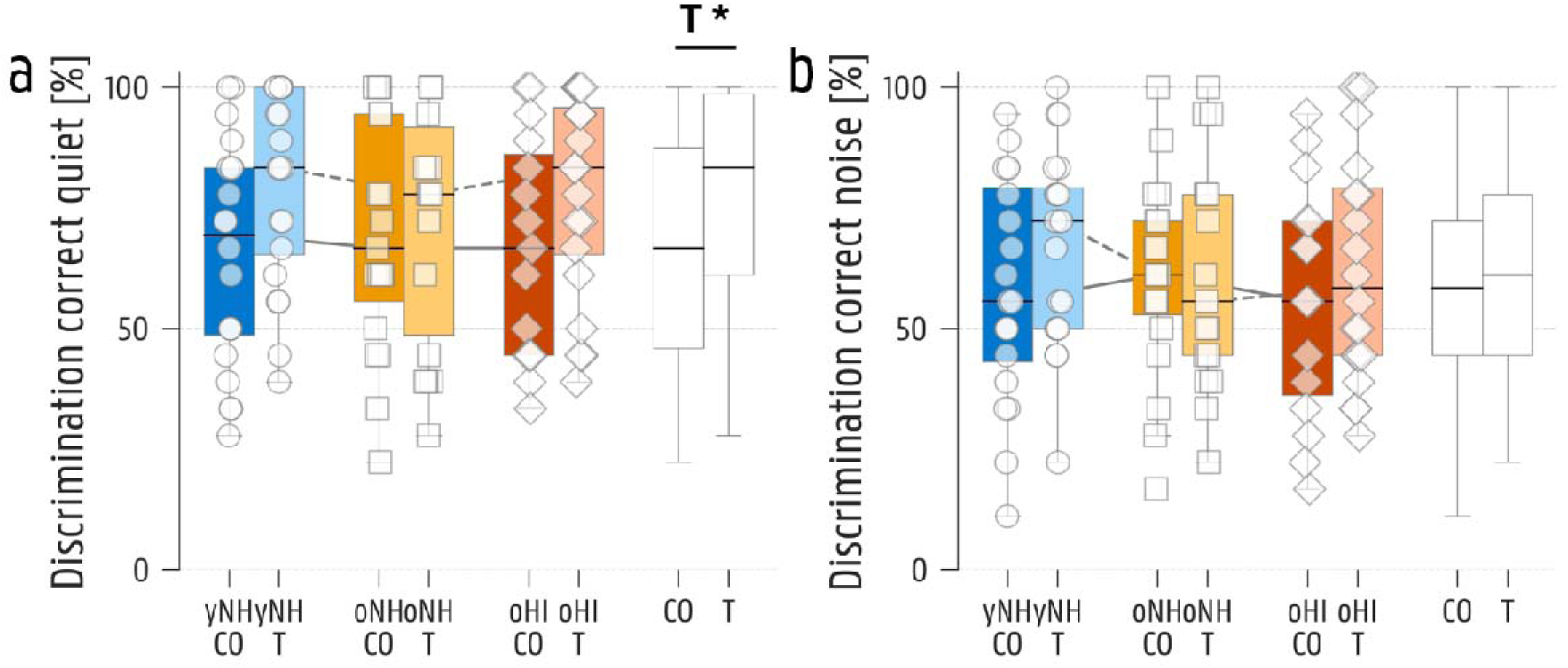
Boxplots of vowel discrimination scores between /o/ and /u/ (a) in quiet and (b) in noise. yNH_CO = young normal hearing control, yNH_T = young normal hearing with tinnitus, oNH_CO = older normal hearing control, oNH_T = older normal hearing with tinnitus, oHI_CO = older hearing impaired control, oHI_T = older hearing impaired with tinnitus, CO = control groups, T = tinnitus groups. *p <.05; **p <.01; ***p <.001.

**Table 5:** Heatmap showing partial eta-squared effect sizes (η²p) for the influence of tinnitus status, age, hearing loss, and their interactions on each dependent variable. Each row represents a measurement outcome, and each column represents a main predictor or interaction term. *p <.05; **p <.01; ***p <.001. T = tinnitus, HL = Hearing Loss, SPIQ = Speech In Quiet, SPIN = Speech In Noise.

|  | Tinnitus | Age | Hearing Loss | T × Age | T × HL |
|---|---|---|---|---|---|
| /O/-/U/ discrimination quiet | * |  |  |  |  |
| /O/-/U/ discrimination noise |  |  |  |  |  |
| SPIQ Low-Pass |  | *** | * |  |  |
| SPIQ High-Pass |  | *** | *** | ** |  |
| SPIN Low-Pass | ** | *** |  |  |  |
| SPIN High-Pass | *** | *** | *** |  |  |

#### 2.2. Matrix sentence test

In the LF SPIQ condition, significant main effects of age (*F*(1,74) = 29.94, *p* <.001, η²⍰ = 0.29; post-hoc yNH_CO vs oNH_CO: *U* = 78.5, *p*_Bonf_ =.004, *r*_rb_ =-0.59, *d* =-0.89; yNH_T vs oNH_T: *U* = 60.0, *p*_Bonf_ <.001, *r_rb_* =-0.68) and hearing loss (*F*(1,69) = 5.86, p =.018, η²⍰ = 0.08; post-hoc oNH_CO vs oHI_CO: *U* = 76.0, *p*_Bonf_ =.044, *r*_rb_ =-0.47) were observed (Table 5). We found no tinnitus-related main or interaction effects (*p* >.05). The SPIQ HF condition also revealed age-related (F(1,74) = 36.95, *p* <.001, η²⍰ = 0.33; post-hoc yNH_T vs oNH_T: *t*(28.7) =-6.51, *p*_Bonf_ <.001, *d* =-2.10) and hearing loss-related main effects (F(1,69) = 66.75, *p* <.001, η²⍰ = 0.49; post-hoc oNH_CO vs oHI_CO: *t*(21.1) =-6.52, *p*_Bonf_ <.001, *d* =-2.32; oNH_T vs oHI_T: *t*(28.3) =-5.35, *p*_Bonf_ <.001, *d* =-1.70). Additionally, a significant Tinnitus x Age interaction emerged (F(1,74) = 7.37, *p* =.008, η²⍰ = 0.09), again driven by a trend toward better performance in the yNH_T group relative to yNH_CO, and poorer performance in the oNH_T group relative to oNH_CO.

Turning to the SPIN condition, the low-pass filtered condition revealed a significant age effect (*F*(1,74) = 11.93, *p* <.001, η²⍰ = 0.14: post-hoc yNH_T vs oNH_T: *t*(37) =-3.50, *p*_Bonf_ =.002, *d* =-1.12; Table 5; Figure 6a). A significant tinnitus-related effect was also observed (F(1,109) = 10.84, *p* =.001, η²⍰ = 0.09; post-hoc oHI_CO vs oHI_T: *t*(33) =-2.59, *p*_Bonf_ =.043, *d* =-0.88), with tinnitus groups performing worse than control groups, an opposite pattern than observed in the vowel discrimination task. In the HF condition, all three main effects were significant (age (*F*(1,74) = 25.07, *p* <.001, η²⍰ = 0.25; post-hoc yNH_CO vs oNH_CO: *U* = 85.5, *p*_Bonf_ =.007, *r*_rb_ =-0.55; yNH_T vs oNH_T: *U* = 74.5, *p*_Bonf_ =.002, *r*_rb_ =-0.61), hearing loss (*F*(1,69) = 25.28, *p* <.001, η²⍰ = 0.27; post-hoc oNH_CO vs oHI_CO: *U* = 32.5, *p*_Bonf_ <.001, *r_rb_*=-0.77; oNH_T vs oHI_T: *U* = 77.5, *p*_Bonf_ =.003, *r*_rb_ =-0.59 and tinnitus (*F*(1,109) = 12.26, *p* <.001, η²⍰ = 0.10; Figure 6b), with tinnitus groups again performing worse than their matched control groups. No significant interactions were observed for SPIN (*p* >.05).

**Figure 6:**
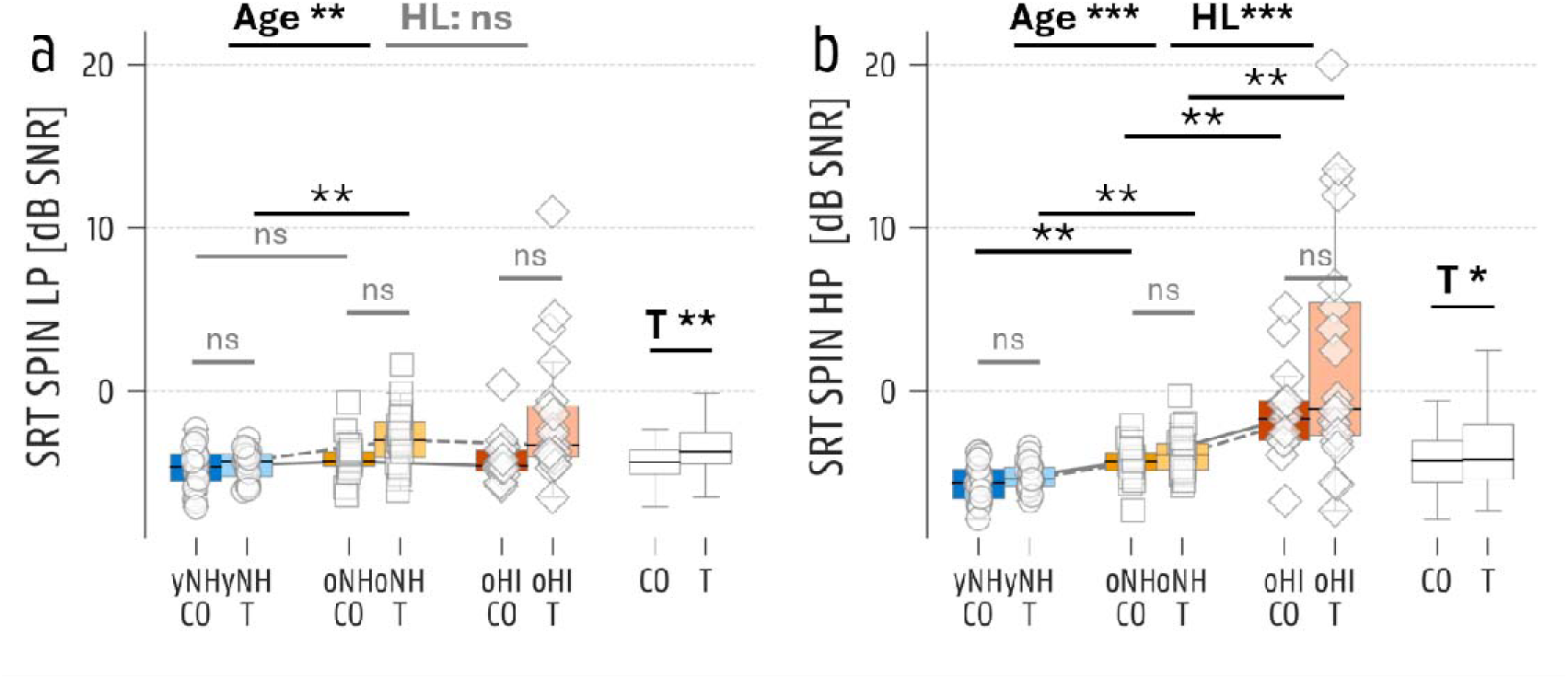
Boxplots of speech in noise reception thresholds (SRT) using the Flemish Matrix test (a) in a low-pass (LP) filtered and (b) in a high-pass (HP) filtered condition. yNH_CO = young normal hearing control, yNH_T = young normal hearing with tinnitus, oNH_CO = older normal hearing control, oNH_T = older normal hearing with tinnitus, oHI_CO = older hearing impaired control, oHI_T = older hearing impaired with tinnitus, CO = control groups, T = tinnitus groups. *p <.05; **p <.01; ***p <.001, ns = not significant.

### 3. Group differences after controlling for OHC damage

Since observed tinnitus-related differences may partly reflect subtle underlying differences in peripheral hearing, all group analysis were repeated using variables residualized against DPOAE_LF and DPOAE_HF amplitudes. As expected, and seen in Table 6, this correction eliminated several of the age-and hearing-related effects observed in the primary analyses. With respect to tinnitus-related ABR amplitude differences, the wave I/V significancy remained (F(1,103) = 4.99, *p* =.028, η²⍰ = 0.05; post-hoc yNH_CO vs yNH_T: *U* = 276.0, *p*_Bonf_ =.017, *r*_rb_ = 0.53), as well as the Tinnitus x Age interactions for wave I/V (F(1,70) = 9.28, *p* =.003, η²⍰ = 0.12), and wave V (F(1,70) = 6.35, *p* =.014, η²⍰ = 0.08). The ABR wave V latency at 70 dB remained significantly different for tinnitus (F(1,104) = 5.50, *p* =.021, η²⍰ = 0.05) and age (F(1,70) = 5.37, *p* =.023, η²⍰ = 0.07). Tinnitus x Age interactions for the EFR remained weakly significant after correction (F(1,70) = 4.54, *p* =.037, η²⍰ = 0.06; post-hoc yNH_T vs oNH_T: *t*(30.2) = 2.70, *p*_Bonf_ =.022, *d* = 0.89), whereas the age-related main differences disappeared (*p* >.05).

**Table 6:** Heatmap for variables residualized against mean low-and high frequency Distortion Product Otoacoustic Emissions (DPOAEs), showing partial eta-squared effect sizes (η²p) for the influence of tinnitus status, age, hearing loss, their interactions, and correlations with low-and high-frequency DPOAEs for each dependent variable. Each row represents a measurement outcome, and each column represents a main predictor or interaction term. *p <.05; **p <.01; ***p <.001. T = tinnitus, HL = Hearing Loss, DPOAE = Distortion Product Otoacoustic Emissions, ABR = Auditory Brainstem Response, amp = amplitude, lat = latency, EFR = Envelope Following Response, FFR = Frequency Following Response, ENV = Envelope, TFS = Temporal Fine Structure, MEMR = Middle Ear Muscle Reflex, MOCR = Medial Olivocochlear Reflex, CEOAE = Click Evoked Otoacoustic Emissions, SPIQ = Speech In Quiet, SPIN = Speech In Noise.

|  | Tinnitus | Age | Hearing Loss | T × Age | T × HL |
|---|---|---|---|---|---|
| DPOAE low frequencies |  | *** | ** |  |  |
| DPOAE high frequencies |  | *** | *** |  |  |
| ABR wave I amp 70 dB peSPL<br>DPGram_LF *r* = 0.19, *p* = .049<br>DPGram_HF *r* = -0.02, *p* = .855 |  |  |  |  |  |
| ABR wave I amp 90 dB peSPL<br>DPGram_LF *r* = 0.26, *p* = .006<br>DPGram_HF *r* = 0.28, *p* = .004 |  | * |  |  |  |
| ABR wave V amp 70 dB peSPL<br>DPGram_LF *r* = 0.31, *p* = .001<br>DPGram_HF *r* = 0.39, *p* < .001 |  |  |  |  |  |
| ABR wave V amp 90 dB peSPL<br>DPGram_LF *r* = 0.33, *p* < .001<br>DPGram_HF *r* = 0.27, *p* = .005 |  |  |  | * |  |
| ABR wave I/V 70 dB peSPL<br>DPGram_LF *r* = -0.17, *p* = .077<br>DPGram_HF *r* = -0.26, *p* = .007 |  |  |  |  |  |
| ABR wave I/V 90 dB peSPL<br>DPGram_LF *r* = 0.02, *p* = .829<br>DPGram_HF *r* = 0.06, *p* = .549 | * |  |  | ** | * |
| ABR wave I lat 70 dB peSPL<br>DPGram_LF *r* = -0.20, *p* = .041<br>DPGram_HF *r* = -0.26, *p* = .008 |  |  |  |  |  |
| ABR wave I lat 90 dB peSPL<br>DPGram_LF *r* = -0.06, *p* = .507<br>DPGram_HF *r* = -0.18, *p* = .061 |  |  |  |  |  |
| ABR wave V lat 70 dB peSPL<br>DPGram_LF *r* = -0.27, *p* = .004<br>DPGram_HF *r* = -0.50, *p* < .001 | * | * |  |  |  |
| ABR wave V lat 90 dB peSPL<br>DPGram_LF *r* = -0.22, *p* = .023<br>DPGram_HF *r* = -0.45, *p* < .001 |  |  |  |  |  |
| /U/-FFR ENV 70 dB SPL<br>DPGram_LF *r* = 0.32, *p* < .001<br>DPGram_HF *r* = 0.35, *p* < .001 |  | * |  |  |  |
| /U/-FFR TFS 70 dB SPL<br>DPGram_LF *r* = 0.43, *p* < .001<br>DPGram_HF *r* = 0.52, *p* < .001 |  | *** |  |  |  |
| EFR 4 kHz 70 dB SPL<br>DPGram_LF *r* = 0.47, *p* < .001<br>DPGram_HF *r* = 0.63, *p* < .001 |  |  |  | * |  |
| MEMR ipsi threshold<br>DPGram_LF *r* = -0.38, *p* < .001<br>DPGram_HF *r* = -0.37, *p* < .001 |  |  |  |  |  |
| MEMR ipsi amplitude at 90 dB sPL<br>DPGram_LF *r* = 0.39, *p* < .001<br>DPGram_HF *r* = 0.41, *p* < .001 |  |  |  |  |  |
| MOCR ipsi ΔCEOAE<br>DPGram_LF *r* = 0.56, *p* < .001<br>DPGram_HF *r* = 0.54, *p* < .001 |  |  |  |  |  |
| LF MOCR ipsi ΔCEOAE<br>DPGram_LF *r* = 0.43, *p* < .001<br>DPGram_HF *r* = 0.38, *p* = .002 |  | * |  |  |  |
| HF MOCR ipsi ΔCEOAE<br>DPGram_LF *r* = 0.45, *p* < .001<br>DPGram_HF *r* = 0.48, *p* < .001 |  |  |  |  |  |
| /O/-/U/ discrimination quiet<br>DPGram_LF *r* = 0.07, *p* = .480<br>DPGram_HF *r* = -0.08, *p* = .430 | * |  |  |  |  |
| /O/-/U/ discrimination noise<br>DPGram_LF *r* = 0.05, *p* = .615<br>DPGram_HF *r* = -0.02, *p* = .797 |  |  |  |  |  |
| SPIQ Low-Pass<br>DPGram_LF *r* = -0.52, *p* < .001<br>DPGram_HF *r* = -0.41, *p* < .001 |  |  |  |  |  |
| SPIQ High-Pass<br>DPGram_LF *r* = -0.62, *p* < .001<br>DPGram_HF *r* = -0.72, *p* < .001 |  |  | *** | * |  |
| SPIN Low-Pass<br>DPGram_LF *r* = -0.19, *p* = .043<br>DPGram_HF *r* = -0.22, *p* = .018 | ** |  |  |  |  |
| SPIN High-Pass<br>DPGram_LF *r* = -0.45, *p* < .001<br>DPGram_HF *r* = -0.44, *p* < .001 |  |  |  |  |  |

For MEMR, all age-related differences were no longer significant after the DPOAE-correction (*p* >.05). For the HF MOCR, the age-related effect disappeared as well, while the LF MOCR comparison shifted from tinnitus-related (*p* <.05) to a significant age-related effect (F(1,45) = 4.77, *p* =.034, η²⍰ = 0.10), suggesting that the initial tinnitus finding may have been confounded by peripheral differences. For the behavioral tasks, tinnitus-related advantage in vowel discrimination remained significant after correction (F(1,106) = 5.84, *p* =.017, η²⍰ = 0.05), as did the tinnitus-related deficit in the SPIN LF task (F(1,107) = 8.12, *p* =.005, η²⍰ = 0.07). SPIQ HF remained significant for hearing loss (F(1,68) = 17.92, *p* <.001, η²⍰ = 0.21; post-hoc oNH_CO vs oHI_CO: *t*(32) =-3.99, *p*_Bonf_ <.001, *d* =-1.38) and Tinnitus x Age interaction (F(1,72) = 4.88, *p* =.030, η²⍰ = 0.06).

### 4. Correlation analyses after controlling for OHC damage

#### 4.1. Relation between objective sensorineural markers

All remaining correlations after correcting for DPOAE amplitudes are visualized in Figure 7. Only one correlation between the objective sensorineural markers (ABR, FFR, EFR, MEMR, MOCR) was present in the control group. ABR wave I correlated with MEMR (ρ(35) =.51, *p* =.001; yNH_CO: r(17) =.58, *p* =.010), a suggested marker for distal auditory nerve encoding.

**Figure 7:**
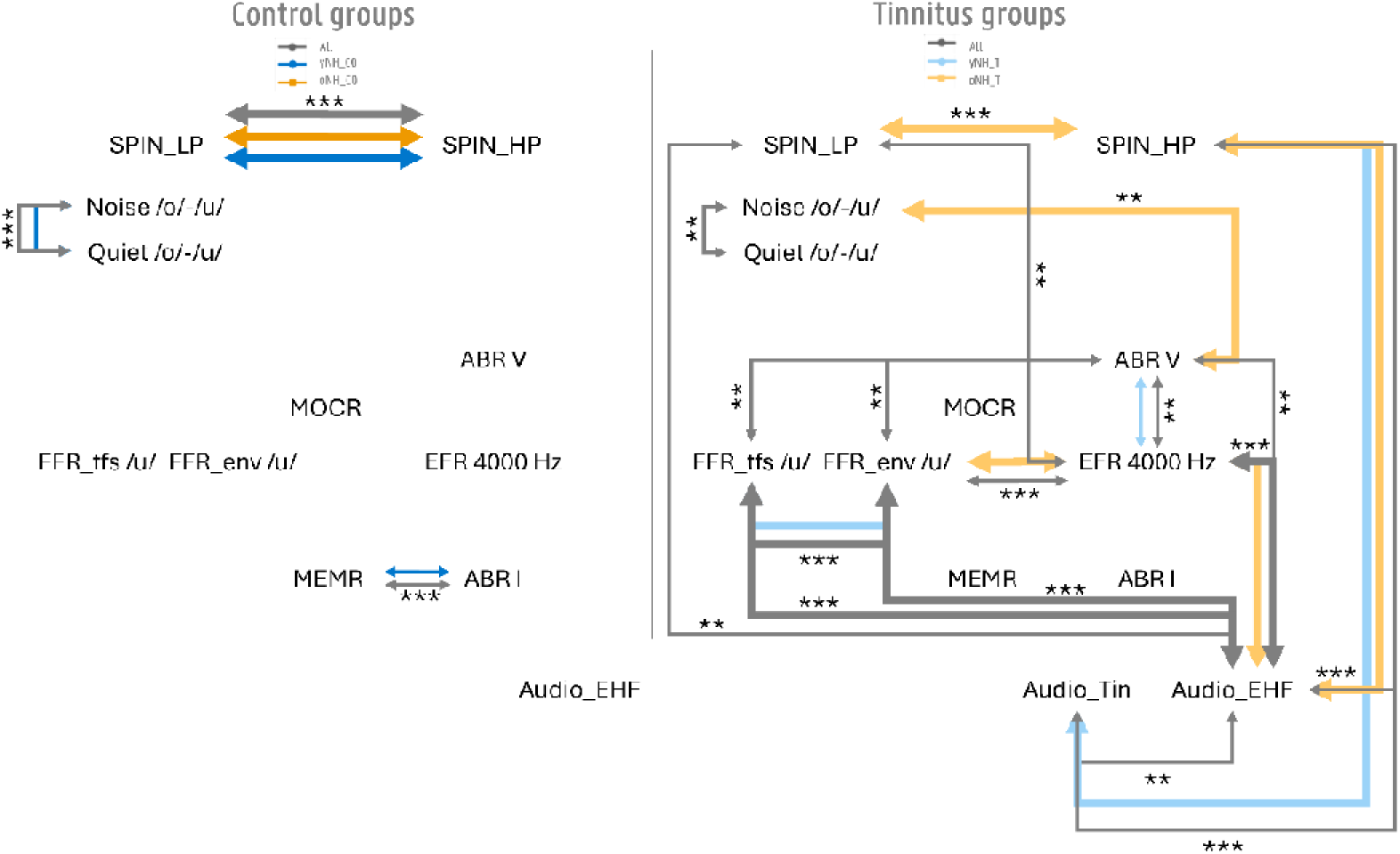
Schematic overview of the significant correlations in the normal hearing groups that still remain after corrections for DPOAE_LF and DPOAE_HF. Grey = young and old groups combined; full arrows: r [0.4 – 0.6]; Thick arrows: r [0.6 – 0.8]; yNH_CO = young normal hearing control, yNH_T = young normal hearing with tinnitus, oNH_CO = older normal hearing control, oNH_T = older normal hearing with tinnitus, oHI_CO = older hearing impaired control, oHI_T = older hearing impaired with tinnitus, CO = control groups, T = tinnitus groups; **p <.01; ***p <.001.

In the tinnitus groups, these distal auditory nerve correlations were absent. Instead, EFR magnitudes correlated with markers of higher brainstem processing (inferior colliculus), namely ABR wave V (r(35) =.49, *p* =.002; yNH_T: r(17) =.60, *p* =.007) and FFR ENV (ρ(35) =.56, *p* <.001; oNH_T: r(16) =.67, *p* =.002). Correspondingly, ABR wave V was significantly correlated with both FFR ENV (ρ(35) =.49, *p* =.002) and FFR TFS (r(35) =.47, *p* =.004). In contrast to the control group, the tinnitus groups showed a strong positive correlation between FFR ENV and TFS magnitudes (ρ(35) =.66, *p* <.001; yNH_T: r(17) =.72, *p* <.001).

Potential interactions with the audiometric thresholds were only observed in the tinnitus groups. Especially EHF audiometry was significantly related to electrophysiological markers EFR (r(35) =-.66, *p* <.001; oNH_T: r(16) =-.63, *p* =.005), ABR wave V amplitude (r(35) =-.49, *p* =.002), and both FFR TFS (r(35) =-.61, *p* <.001) and FFR ENV (ρ(35) =-.64, *p* <.001). Also SPIN was related to EHF audiometry for both LF (ρ (35) =.50, p =.002) and HF conditions (r(35) =.52, *p* <.001; yNH_T: r(17) =.82, *p* <.001; oNH_T: r(16) =.72, *p* <.001). Audiometry thresholds at the tinnitus pitch (Audio_Tin) were only associated with HF SPIN (ρ(35) =.54, *p* <.001; yNH_T: r(17) =.70, *p* <.001) and EHF audiometry (ρ(35) =.48, *p* =.003).

#### 4.2. Relation between sensorineural markers and functional speech-related outcomes

In addition to the correlations among objective neural markers, Figure 7 shows associations between objective markers and speech-related task performance in the four groups with normal audiometric thresholds. The /o/-/u/ vowel discrimination task, which showed a trend toward improved performance in the tinnitus group, yielded only one significant correlation, with ABR wave V in the oNH_T group (r(15) =.64, *p* =.006). Between the quiet and noise condition, the vowel discrimination scores were correlated in both the tinnitus (ρ(34) =.52, *p* =.001) and control groups (r(37) =.54, *p* <.001; yNH_CO: r(18) =.60, *p* =.006).

In the tinnitus groups, LF SPIN correlated with EFR magnitude (ρ(35) =-.50, *p* =.002). No other significant correlations could be observed between SPIN and the objective sensorineural markers. Between LF and HF SPIN, SRTs correlated strongly in the control group (r(37) =.74, *p* <.001; yNH_CO: r(18) =.80, *p* <.001; oNH_CO: r(17) =.67, *p* =.002), but not in the tinnitus groups *(p* >.05). This may suggest that control participants rely on similar encoding mechanisms for both LF and HF SPIN perception, whereas tinnitus participants may additionally rely on other processing strategies, potentially related to ENV encoding, as reflected in the EFR–LF SPIN correlation (Figure 7).

Audiometric thresholds only showed a significant correlation between EHF audiometry and TFI scores, with higher TFI scores being related to worse hearing thresholds (r(35) =.47, *p* =.004).

#### 4.3. Relation with subjective tinnitus-related parameters

Finally, we examined correlations between the objective markers and subjective tinnitus-and hyperacusis-related parameters (Figure 8). Tinnitus distress was assessed using the TFI, and tinnitus psychoacoustics were characterized using tinnitus loudness and minimal masking levels. Strong correlations were found between higher perceived tinnitus loudness and worse HF SPIN performance (r(35) =.64, *p* <.001; yNH_T: r(17) =.64, *p* =.003; oNH_T: r(16) =.69, *p* =.001). None of the objective markers correlated significantly to TFI or minimal masking levels; however, TFI and minimal masking levels were significantly correlated with each other in the yNH_T group, with higher tinnitus distress (higher TFI scores) related to lower masking thresholds (r(17) =-.66, *p* =.002).

**Figure 8:**
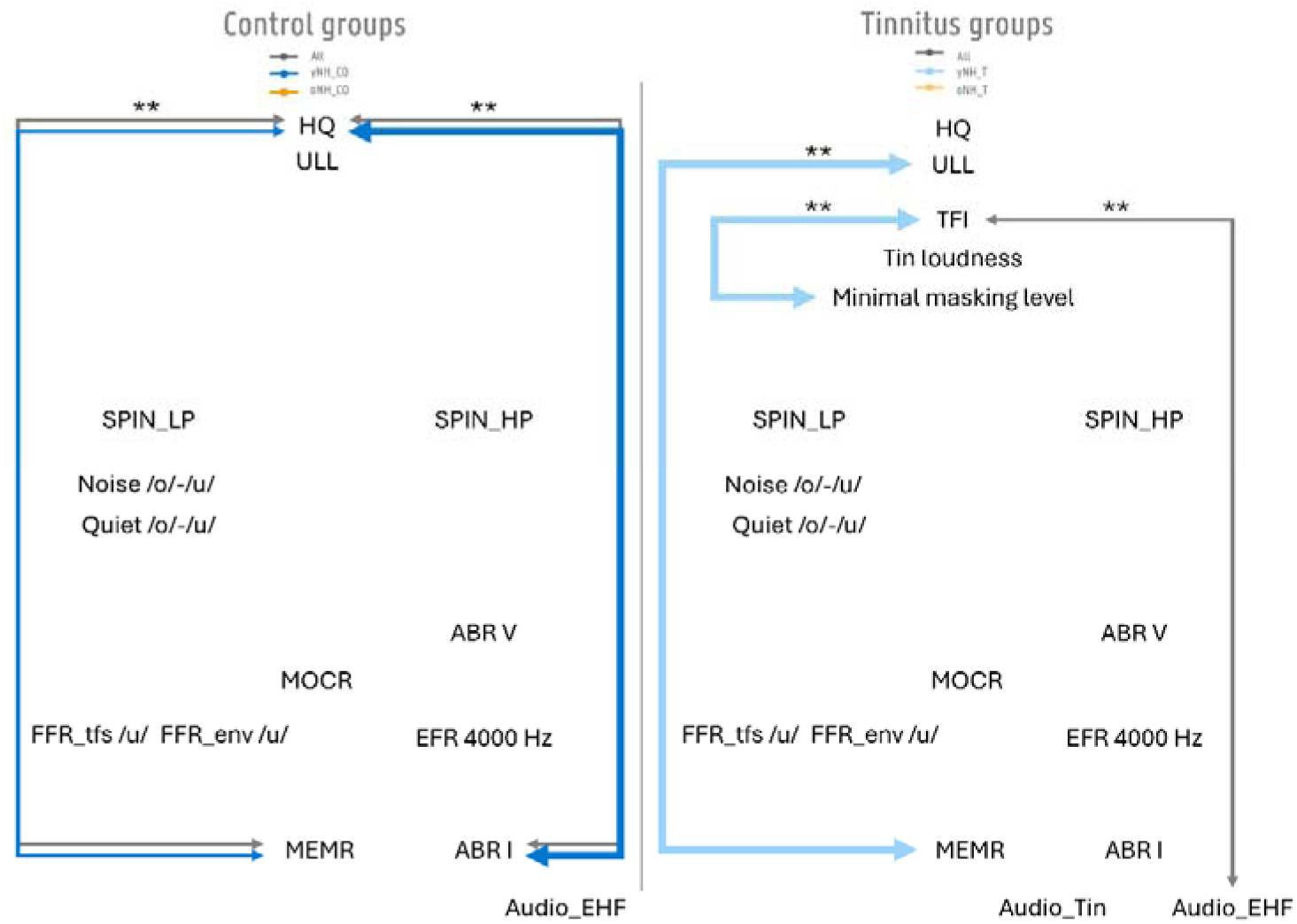
Schematic overview of the significant correlations with tinnitus-and hyperacusis-related parameters in the normal hearing groups that still remain after corrections for DPOAE_LF and DPOAE_HF. Grey = young and old groups combined; full arrows: r [0.4 – 0.6]; Thick arrows: r [0.6 – 0.8]; yNH_CO = young normal hearing control, yNH_T = young normal hearing with tinnitus, oNH_CO = older normal hearing control, oNH_T = older normal hearing with tinnitus, oHI_CO = older hearing impaired control, oHI_T = older hearing impaired with tinnitus, CO = control groups, T = tinnitus groups; **p <.01; ***p <.001.

Although participants with hyperacusis were excluded based on HQ scores, residual variability in both HQ and ULL measures was observed, which correlated with peripheral sensorineural markers. Correlations were observed with MEMR amplitude within both groups. In the tinnitus group, MEMR correlated strongly with ULL (yNH_T: r(17) =.60, *p* =.006), indicating that smaller MEMR responses are related to lower ULLs, and reduced sound-level tolerance. Similar trends were observed in the control groups: higher HQ scores were associated with lower MEMR amplitudes (ρ(37) =-.46, *p* =.004; yNH_CO: r(18) =-.56, *p* =.010). In this group, ABR wave I amplitude also correlated with HQ scores (r(35) =-.47, *p* =.003; yNH_CO: r(17) =-.67, *p* =.002), consistent with the previously noted correlation between ABR wave I and MEMR amplitude in this group.

## Discussion

### 1. Central brainstem-related enhancement was only detectable in young subjects with tinnitus

Across nearly all objective markers of afferent sensorineural encoding, we observed clear age-and hearing-loss-related effects. Tinnitus-related effects were less pronounced and appeared mainly in the young groups, as evidenced by Tinnitus x Age interactions showing a tendency toward enhanced ABR wave V amplitudes, greater central gain based on ABR I/V ratios, and elevated EFR responses. However, such effects had small effect sizes, especially after DPOAE correction. Unlike several previous studies reporting reduced wave I amplitudes (Bramhall, Konrad-Martin, & McMillan, 2018; Bramhall et al., 2019; Möhrle et al., 2019; Morse & Vander Werff, 2023), or enlarged V/I ratios (Sendesen et al., 2022; Song et al., 2018), the present study revealed a tendency toward enhanced wave V responses, rather than wave I attenuation. Following DPOAE-corrections, wave I amplitudes remained associated with age, whereas wave V did not, suggesting that wave I changes should be interpreted with caution given potential age-related confounds. Despite these discrepancies, the overall pattern is compatible with central gain theories, indicative of compensatory gain along the brainstem, but remains tentative given the small effects, the dependence of the ratio on wave V, and the non-significant within-group wave V difference. Beyond age-related differences, cross-study inconsistencies may stem from methodological variation, including differences in stimulus intensity, sex, or amplitude-calculation methods.

Potential EFR enhancements in individuals with tinnitus have not been consistently reported elsewhere. Guest et al. (2017) found no EFR or ABR wave I differences in young listeners. The primary difference between that study and the present, is that it employed a sinusoidal rather than rectangular amplitude modulation, which may have reduced sensitivity to cochlear synaptopathy (Vasilkov et al., 2021). Similarly, Devolder et al. (2024) reported no EFR differences. This absence of enhancements may be attributable to lower tinnitus distress (based on TFI) in that cohort, as well as a higher proportion of participants reporting hyperacusis compared to the present study. Nevertheless, the significance of EFR-related enhancements in the present study was only weakly significant after DPOAE-correction.

In the control groups, ABR wave I and MEMR amplitudes, both indexing distal sensorineural hearing, correlated with one another after DPOAE correction. Surprisingly, distal correlations were absent in the tinnitus groups. This suggests that tinnitus-related central gain or neural noise may alter these measures. Across DPOAEs, ABR, EFR, and FFR measures, we found no evidence of increased hidden sensorineural hearing loss in the chronic tinnitus groups. Sensorineural hearing deficits may be transient in the acute stage or may require frequency-specific testing at the tinnitus pitch. Lefeuvre et al. (2019) note that tinnitus-related hearing loss may go undetected even on the tonal audiogram, owing to its limited frequency specificity. Future studies should examine potential effects of central or brainstem gain on EFR and other FFR parameters, characterize sensorineural hearing in the acute tinnitus stage, and evaluate the use of eliciting stimuli centered on the tinnitus frequency.

### 2. While vowel discrimination is improved in subjects with tinnitus, functional speech-in-noise intelligibility is impaired

While the current study demonstrated enhanced performance on a basic psychoacoustic task in individuals with tinnitus, their functional SPIN performance was impaired. This discrepancy between tasks is consistent with the broader tinnitus literature, where outcomes vary widely depending on stimulus characteristics, noise type, task complexity, and cognitive demands. This pattern aligns with findings from Jagoda et al. (2018), who reported that higher tinnitus distress was associated with poorer SPIN performance but better vowel discrimination, with longer reaction times. Their electrophysiological N2 oddball data suggested heightened sensitivity to salient auditory input, possibly reflecting insufficient inhibition of background noise. Such hyperresponsiveness may hinder speech perception under challenging listening conditions.

Listening effort may further explain the divergence between vowel discrimination and sentence-based SPIN (Degeest, Keppler, & Corthals, 2017; Degeest, Kestens, & Keppler, 2022). Although both tasks require attention, the sentence test requires greater cognitive load, as participants must retain the entire sentence in working memory. This could disproportionately affect older participants who are more vulnerable to cognitive difficulties (Degeest, Keppler, & Corthals, 2015). This also accords with our observation that tinnitus loudness correlated with SPIN performance, suggesting that louder tinnitus may serve as a distraction during the SPIN task. Sevmez and Sendesen (2026) recently showed a relation between tinnitus and reduced cognitive learning and memory. Nevertheless, this was not correlated with SPIN performance in their study. Future studies should account the potential link between cognitive function and speech perception in individuals with tinnitus.

Another plausible explanation relates to temporal processing. Tinnitus-related SPIN differences were more pronounced in the low-pass SPIN condition, especially after DPOAE-correction. Moreover, in the tinnitus groups, EFRs correlated with low-pass SPIN scores, whereas in the controls, no sensorineural markers correlated with high-frequency SPIN. Furthermore, the vowel discrimination task employed the vowels /o/ and /u/, which differ exclusively in their low-frequency spectral content. Low-frequency sounds provide access to both temporal ENV and TFS cues, whereas high-frequency perception primarily relies on ENV due to reduced phase locking. Collectively, these findings suggest that tinnitus could be related to alterations in temporal processing, as proposed in earlier research (Auerbach, Rodrigues, & Salvi, 2014; Bureš et al., 2019; Devolder et al., 2024).

The absence of correlations between the SPIN LF and HF condition in the younger tinnitus group may reflect distinct underlying auditory encoding mechanisms in contrast to the control groups, where LF and HF conditions correlated strongly. The observed association between LF SPIN and EFR, together with the correlations among markers of the higher brainstem (inferior colliculus level; EFR with ABR V and FFR), may indicate a reliance on ENV encoding, rather than TFS. Sevmez and Sendesen (2026) examined TFS sensitivity in subjects with tinnitus. They found reduced TFS sensitivity and SPIN performance using the Matrix sentence test, but this was not related to each other. Nevertheless, the compared test groups differed based on extended high frequency thresholds and age. Indeed, Ding et al. (2022) previously administrated the same TFS test in participants with tinnitus but observed no group differences, instead reporting a significant correlation with age and extended high frequency thresholds. To date, no evidence of reduced TFS sensitivity has been established; the FFR data in the present study likewise yielded no associations with TFS processing. Some studies, for example Goossens et al. (2018) and Millman et al. (2017) suggest that enhanced envelope encoding, which may be present in tinnitus based on our EFR outcomes, can negatively impact speech comprehension. It would also be relevant to investigate whether these low-frequency behavioral differences are related to the observed shift towards reduced low-frequency MOCR activity in patients with tinnitus. Future work should disentangle low-versus high-frequency processing differences in tinnitus using psychoacoustic or objective ENV-and TFS-specific measures in age-stratified test groups.

### 3. The MEMR showed no tinnitus-related differences, but can potentially be related to hyperacusis complaints

A final aspect that emerged from the correlation analysis was an association between MEMR and hyperacusis-related parameters, specifically HQ scores and ULLs. Prior MEMR studies in tinnitus (Guest, Munro, & Plack, 2019; Vasilkov et al., 2023) have not yet systematically included hyperacusis measures. One possible explanation for this association is peripheral hidden hearing loss. We also observed correlations between HQ scores and ABR wave I amplitudes, suggesting reduced auditory nerve output in individuals reporting more hyperacusis symptoms. This aligns with the framework of Zeng (2020), which proposes that tinnitus may arise from neural noise triggered by hearing loss, whereas hyperacusis may be more strongly linked to central gain mechanisms following hidden hearing loss.

Another possibility is that impaired stapedial reflex function reduces the ear’s natural attenuation of intense stimuli, making everyday sounds seem uncomfortably loud to affected listeners. This potential relationship between auditory reflexes and hyperacusis has also been suggested by Gordon (1986) and Saxena et al. (2020), and has been documented in patients with Williams syndrome (Attias et al., 2008; Gothelf et al., 2006; Silva et al., 2021). Because the current study intentionally excluded participants with hyperacusis complaints, these associations should be interpreted cautiously. Future studies should explicitly include individuals with hyperacusis complaints to further clarify the relationship between MEMR function, auditory nerve integrity, central gain, and sound-level tolerance.

### 4. Conclusion

This study demonstrated that afferent sensorineural auditory processing is strongly shaped by age and hearing loss, whereas tinnitus-related effects on sensorineural markers are weaker and age-dependent. In young normal-hearing listeners, tinnitus-related differences were reflected in enhanced ABR wave V amplitudes, elevated EFR responses, and improved vowel discrimination, suggesting altered central auditory weighting without measurable audiometric-or OHC-related impairment. In contrast, older listeners with tinnitus showed no neural alterations and poorer speech-in-noise performance, indicating that tinnitus-related changes may interact with age-related sensorineural decline. Although efferent reflex measurements (MOCR and MEMR) did not consistently differ between groups, MEMR was potentially associated with hyperacusis-related parameters. This relationship warrants further investigation, particularly regarding speech encoding and central gain in individuals with hyperacusis.

Overall, these findings point to distinct tinnitus-related profiles across the lifespan: central enhancement and preserved perception in younger adults versus general reduced sensorineural encoding and increased speech understanding difficulties in older adults. This underscores the need for age-specific interpretation of auditory sensorineural biomarkers in tinnitus-related research, with particular attention to possible central gain effects that may confound relationships among auditory measurements.

The authors declare no competing financial interests.

## Acknowledgments

We thank François Deloche and Attila Fráter for assistance with the measurement setup and calibration, and all participants for their time and commitment to this study. This work was supported by the Fonds Wetenschappelijk Onderzoek (FWO), grant G068621N (AuDiMod).

## Supplementary

**Table S1:** Descriptive statistics (mean, standard deviation, and sample size) for all outcome measures across the six test groups.

|  | **yNH_CO** | **yNH_T** | **oNH_CO** | **oNH_T** | **oHI_CO** | **oHI_T** |
|---|---|---|---|---|---|---|
| **Audiogram**<br>**low frequencies** | M = 1.125<br>SD = 2.807<br>n = 20 | M = -0.375<br>SD = 2.927<br>n = 20 | M = 4.276<br>SD = 4.296<br>n = 19 | M = 5.329<br>SD = 4.406<br>n = 19 | M = 11.000<br>SD = 5.004<br>n = 15 | M = 11.312<br>SD = 8.473<br>n = 20 |
| **Audiogram**<br>**high frequencies** | M = 5.000<br>SD = 5.419<br>n = 20 | M = 2.400<br>SD = 4.333<br>n = 20 | M = 9.053<br>SD = 5.126<br>n = 19 | M = 13.947<br>SD = 5.977<br>n = 19 | M = 35.867<br>SD = 10.246<br>n = 15 | M = 38.050<br>SD = 12.804<br>n = 20 |
| **Audiogram**<br>**extended high frequencies** | M = 11.500<br>SD = 13.381<br>n = 20 | M = 8.188<br>SD = 7.890<br>n = 20 | M = 39.079<br>SD = 11.806<br>n = 19 | M = 48.421<br>SD = 11.821<br>n = 19 | M = 65.583<br>SD = 11.229<br>n = 15 | M = 66.125<br>SD = 9.867<br>n = 20 |
| **DPOAE**<br>**low frequencies** | M = 8.114<br>SD = 5.036<br>n = 20 | M = 7.064<br>SD = 4.467<br>n = 19 | M = 4.009<br>SD = 5.070<br>n = 19 | M = 3.266<br>SD = 4.738<br>n = 18 | M = 1.462<br>SD = 6.061<br>n = 15 | M = -1.368<br>SD = 5.543<br>n = 20 |
| **DPOAE**<br>**high frequencies** | M = 1.448<br>SD = 4.810<br>n = 20 | M = 0.059<br>SD = 4.678<br>n = 19 | M = -2.467<br>SD = 3.390<br>n = 19 | M = -3.884<br>SD = 4.244<br>n = 18 | M = -8.610<br>SD = 3.619<br>n = 15 | M = -9.046<br>SD = 3.210<br>n = 20 |
| **ABR wave I amp**<br>**70 dB peSPL** | M = 0.136<br>SD = 0.067<br>n = 19 | M = 0.140<br>SD = 0.072<br>n = 20 | M = 0.112<br>SD = 0.124<br>n = 18 | M = 0.121<br>SD = 0.095<br>n = 19 | M = 0.147<br>SD = 0.144<br>n = 15 | M = 0.121<br>SD = 0.106<br>n = 19 |
| **ABR wave I amp**<br>**90 dB peSPL** | M = 0.174<br>SD = 0.083<br>n = 19 | M = 0.145<br>SD = 0.084<br>n = 20 | M = 0.100<br>SD = 0.069<br>n = 18 | M = 0.107<br>SD = 0.065<br>n = 19 | M = 0.134<br>SD = 0.135<br>n = 15 | M = 0.105<br>SD = 0.072<br>n = 18 |
| **ABR wave V amp**<br>**70 dB peSPL** | M = 0.259<br>SD = 0.106<br>n = 19 | M = 0.345<br>SD = 0.151<br>n = 20 | M = 0.215<br>SD = 0.147<br>n = 18 | M = 0.192<br>SD = 0.089<br>n = 19 | M = 0.209<br>SD = 0.157<br>n = 15 | M = 0.160<br>SD = 0.120<br>n = 19 |
| **ABR wave V amp**<br>**90 dB peSPL** | M = 0.255<br>SD = 0.142<br>n = 19 | M = 0.342<br>SD = 0.113<br>n = 20 | M = 0.243<br>SD = 0.107<br>n = 18 | M = 0.197<br>SD = 0.077<br>n = 19 | M = 0.217<br>SD = 0.158<br>n = 15 | M = 0.218<br>SD = 0.123<br>n = 18 |
| **ABR wave I/V**<br>**70 dB peSPL** | M = 0.682<br>SD = 0.567<br>n = 19 | M = 0.470<br>SD = 0.303<br>n = 20 | M = 1.444<br>SD = 2.614<br>n = 18 | M = 0.568<br>SD = 0.551<br>n = 19 | M = 1.277<br>SD = 2.301<br>n = 15 | M = 2.845<br>SD = 4.144<br>n = 19 |
| **ABR wave I/V**<br>**90 dB peSPL** | M = 0.868<br>SD = 0.552<br>n = 19 | M = 0.431<br>SD = 0.224<br>n = 20 | M = 0.472<br>SD = 0.321<br>n = 18 | M = 0.610<br>SD = 0.330<br>n = 19 | M = 1.002<br>SD = 1.137<br>n = 15 | M = 0.457<br>SD = 0.790<br>n = 18 |
| **ABR wave I lat**<br>**70 dB peSPL** | M = 3.984<br>SD = 0.709<br>n = 19 | M = 3.790<br>SD = 0.186<br>n = 20 | M = 4.000<br>SD = 0.837<br>n = 18 | M = 3.984<br>SD = 0.742<br>n = 19 | M = 4.053<br>SD = 0.890<br>n = 15 | M = 4.347<br>SD = 1.145<br>n = 19 |
| **ABR wave I lat**<br>**90 dB peSPL** | M = 8.026<br>SD = 0.679<br>n = 19 | M = 7.740<br>SD = 0.362<br>n = 20 | M = 8.472<br>SD = 0.474<br>n = 18 | M = 8.495<br>SD = 0.869<br>n = 19 | M = 9.273<br>SD = 1.079<br>n = 15 | M = 8.700<br>SD = 1.216<br>n = 19 |
| **ABR wave V lat**<br>**70 dB peSPL** | M = 3.253<br>SD = 0.445<br>n = 19 | M = 3.010<br>SD = 0.168<br>n = 20 | M = 3.228<br>SD = 0.611<br>n = 18 | M = 3.247<br>SD = 0.409<br>n = 19 | M = 3.227<br>SD = 0.623<br>n = 15 | M = 3.267<br>SD = 0.944<br>n = 18 |
| **ABR wave V lat**<br>**90 dB peSPL** | M = 7.126<br>SD = 0.275<br>n = 19 | M = 7.005<br>SD = 0.256<br>n = 20 | M = 7.394<br>SD = 0.501<br>n = 18 | M = 7.416<br>SD = 0.526<br>n = 19 | M = 7.747<br>SD = 0.791<br>n = 15 | M = 7.783<br>SD = 1.075<br>n = 18 |
| **/U/-FFR ENV**<br>**70 dB SPL** | M = 0.063<br>SD = 0.027<br>n = 19 | M = 0.082<br>SD = 0.035<br>n = 20 | M = 0.049<br>SD = 0.020<br>n = 18 | M = 0.044<br>SD = 0.014<br>n = 19 | M = 0.048<br>SD = 0.023<br>n = 15 | M = 0.040<br>SD = 0.031<br>n = 20 |
| **/U/-FFR TFS**<br>**70 dB SPL** | M = 0.082<br>SD = 0.028<br>n = 19 | M = 0.088<br>SD = 0.025<br>n = 20 | M = 0.050<br>SD = 0.019<br>n = 18 | M = 0.050<br>SD = 0.022<br>n = 19 | M = 0.041<br>SD = 0.020<br>n = 15 | M = 0.045<br>SD = 0.021<br>n = 20 |
| **EFR 4 kHz**<br>**70 dB SPL** | M = 0.174<br>SD = 0.124<br>n = 19 | M = 0.212<br>SD = 0.075<br>n = 20 | M = 0.125<br>SD = 0.054<br>n = 18 | M = 0.100<br>SD = 0.069<br>n = 19 | M = 0.072<br>SD = 0.031<br>n = 15 | M = 0.056<br>SD = 0.043<br>n = 20 |
| **MEMR ipsi**<br>**threshold** | M = 70.692<br>SD = 9.534<br>n = 19 | M = 69.872<br>SD = 11.867<br>n = 19 | M = 75.103<br>SD = 6.979<br>n = 15 | M = 76.431<br>SD = 11.186<br>n = 16 | M = 77.150<br>SD = 10.025<br>n = 15 | M = 79.410<br>SD = 10.323<br>n = 17 |
| **MEMR ipsi**<br>**amplitude at 90 dB SPL** | M = 0.238<br>SD = 0.194<br>n = 20 | M = 0.215<br>SD = 0.163<br>n = 20 | M = 0.104<br>SD = 0.084<br>n = 19 | M = 0.122<br>SD = 0.150<br>n = 19 | M = 0.130<br>SD = 0.168<br>n = 15 | M = 0.063<br>SD = 0.087<br>n = 20 |
| **MOCR ipsi ∆CEOAE** | M = 19.488<br>SD = 9.289<br>n = 15 | M = 13.618<br>SD = 11.915<br>n = 9 | M = 17.149<br>SD = 8.368<br>n = 14 | M = 14.712<br>SD = 7.506<br>n = 13 | M = 13.910<br>SD = 5.844<br>n = 5 | M = 7.575<br>SD = 13.129<br>n = 10 |
| **Low frequencies**<br>**MOCR ipsi ∆CEOAE** | M = 18.244<br>SD = 12.774<br>n = 15 | M = 9.759<br>SD = 14.416<br>n = 9 | M = 21.847<br>SD = 12.186<br>n = 14 | M = 14.603<br>SD = 9.954<br>n = 13 | M = 15.345<br>SD = 5.679<br>n = 5 | M = 9.430<br>SD = 17.058<br>n = 10 |
| **High frequencies**<br>**MOCR ipsi ∆CEOAE** | M = 21.901<br>SD = 14.284<br>n = 15 | M = 17.491<br>SD = 8.630<br>n = 9 | M = 12.377<br>SD = 14.428<br>n = 14 | M = 13.236<br>SD = 7.196<br>n = 13 | M = 14.473<br>SD = 6.936<br>n = 5 | M = 0.663<br>SD = 14.566<br>n = 10 |
| **/O/-/U/ discrimination**<br>**quiet** | M = 0.656<br>SD = 0.233<br>n = 20 | M = 0.797<br>SD = 0.202<br>n = 20 | M = 0.696<br>SD = 0.242<br>n = 19 | M = 0.722<br>SD = 0.246<br>n = 18 | M = 0.667<br>SD = 0.233<br>n = 15 | M = 0.783<br>SD = 0.211<br>n = 20 |
| **/O/-/U/ discrimination**<br>**noise** | M = 0.578<br>SD = 0.233<br>n = 20 | M = 0.672<br>SD = 0.201<br>n = 20 | M = 0.602<br>SD = 0.204<br>n = 19 | M = 0.605<br>SD = 0.228<br>n = 18 | M = 0.559<br>SD = 0.246<br>n = 15 | M = 0.628<br>SD = 0.241<br>n = 20 |
| **SPIQ**<br>**Low-Pass** | M = 15.290<br>SD = 4.466<br>n = 20 | M = 16.370<br>SD = 3.786<br>n = 20 | M = 19.995<br>SD = 4.591<br>n = 19 | M = 22.889<br>SD = 5.599<br>n = 19 | M = 26.667<br>SD = 8.639<br>n = 15 | M = 28.335<br>SD = 14.307<br>n = 20 |
| **SPIQ**<br>**High-Pass** | M = 28.335<br>SD = 5.568<br>n = 20 | M = 26.920<br>SD = 3.575<br>n = 20 | M = 32.105<br>SD = 5.729<br>n = 19 | M = 37.458<br>SD = 6.133<br>n = 19 | M = 51.047<br>SD = 10.025<br>n = 15 | M = 53.930<br>SD = 12.238<br>n = 20 |
| **SPIN**<br>**Low-Pass** | M = -4.705<br>SD = 1.348<br>n = 20 | M = -4.510<br>SD = 0.965<br>n = 20 | M = -4.121<br>SD = 1.221<br>n = 19 | M = -2.858<br>SD = 1.824<br>n = 19 | M = -4.167<br>SD = 1.515<br>n = 15 | M = -1.645<br>SD = 4.123<br>n = 20 |
| **SPIN**<br>**High-Pass** | M = -5.615<br>SD = 1.217<br>n = 20 | M = -5.190<br>SD = 0.874<br>n = 20 | M = -4.363<br>SD = 1.154<br>n = 19 | M = -3.832<br>SD = 1.363<br>n = 19 | M = -1.367<br>SD = 2.941<br>n = 15 | M = 1.995<br>SD = 7.499<br>n = 20 |

**yNH\_CO = young normal hearing control, yNH\_T = young normal hearing with tinnitus, oNH\_CO = older normal hearing control, oNH\_T = older normal hearing with tinnitus, oHI\_CO = older hearing impaired control, oHI\_T = older hearing impaired with tinnitus, DPOAE = Distortion Product Otoacoustic Emissions, ABR = Auditory Brainstem Response, amp = amplitude, lat = latency, EFR = Envelope Following Response, FFR = Frequency Following Response, ENV = Envelope, TFS = Temporal Fine Structure, MEMR = Middle Ear Muscle Reflex, MOCR = Medial Olivocochlear Reflex, CEOAE = Click Evoked Otoacoustic Emissions, SPIQ = Speech In Quiet, SPIN = Speech In Noise.**

---

**Figure S1:**
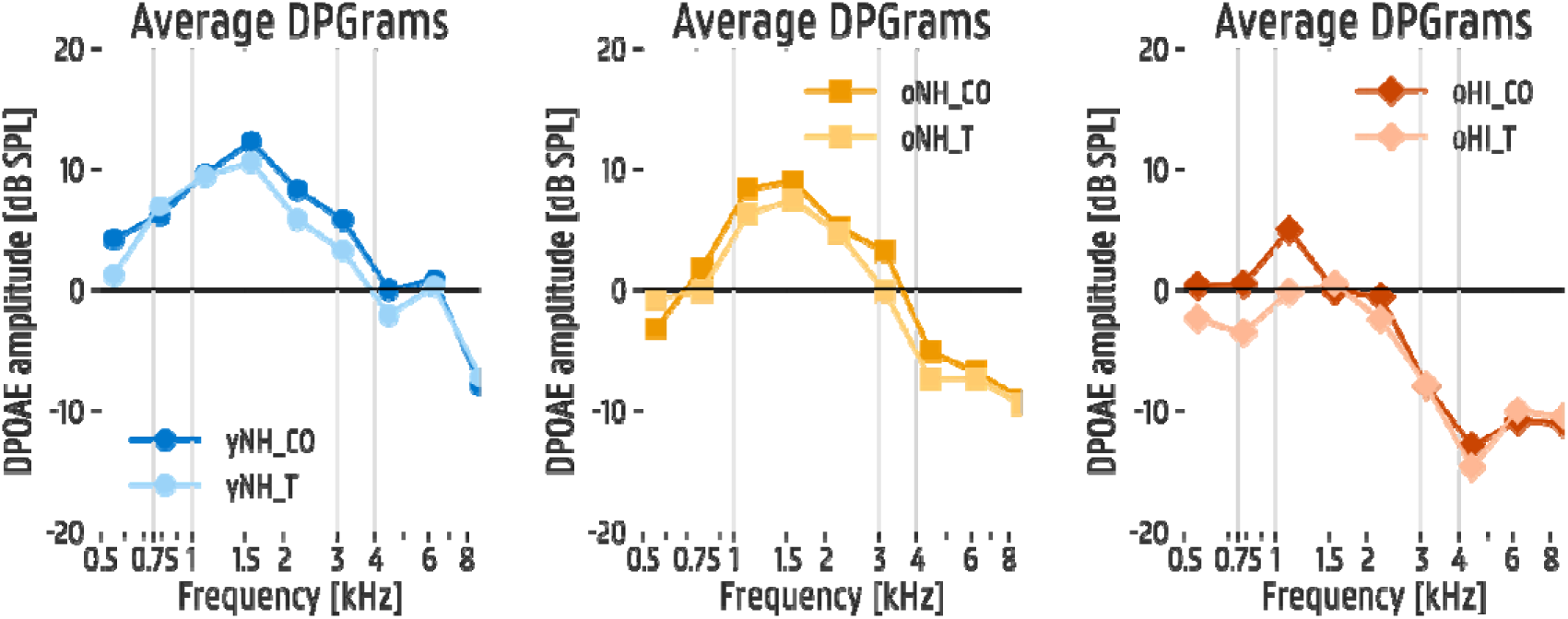
Average distortion product otoacoustic emissions (DPOAE) amplitudes for the six test groups; yNH_CO = young normal hearing control, yNH_T = young normal hearing with tinnitus, oNH_CO = older normal hearing control, oNH_T = older normal hearing with tinnitus, oHI_CO = older hearing impaired control, oHI_T = older hearing impaired with tinnitus.

**Figure S2:**
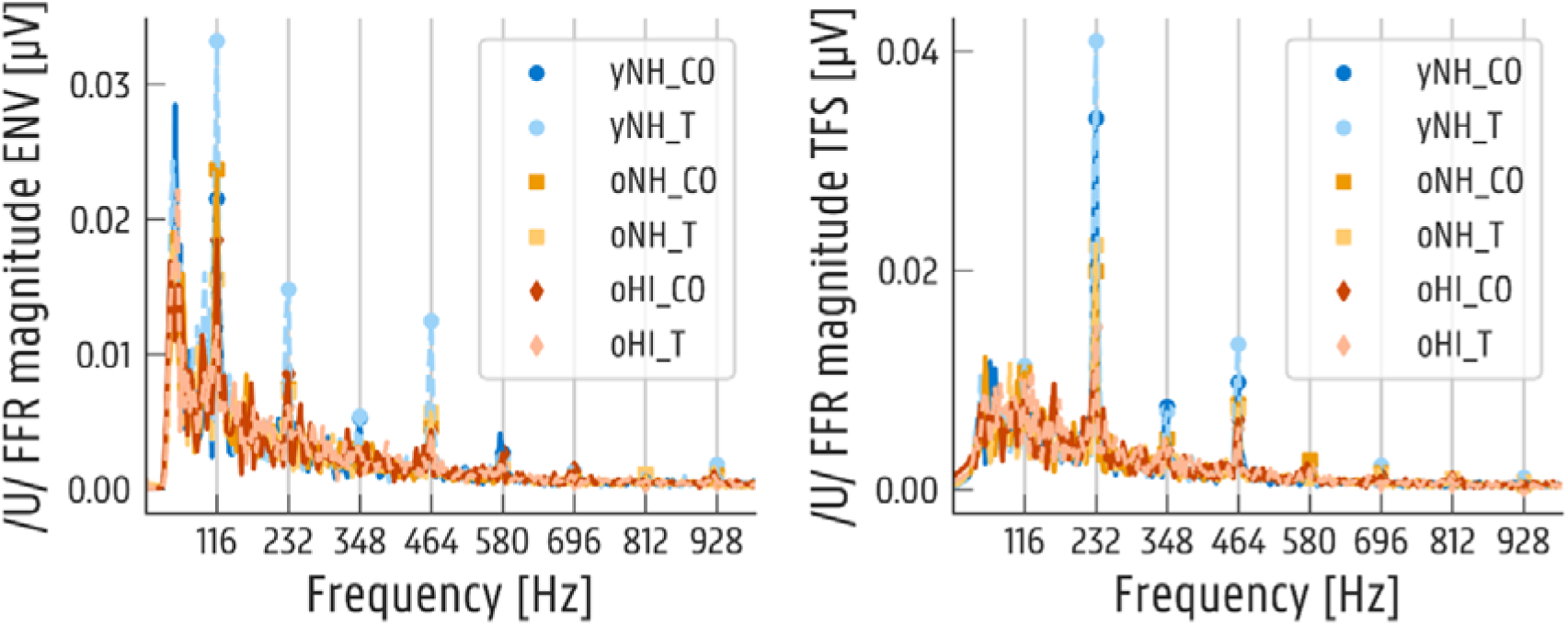
Frequency following responses (FFRs) with a 70 dB SPL /u/-sound stimulus, splitted infot temporal envelope (ENV) and temporal fine structure encoding (TFS); mean FFR responsed per test group with indication of the 116 Hz formant peaks. yNH_CO = young normal hearing control, yNH_T = young normal hearing with tinnitus, oNH_CO = older normal hearing control, oNH_T = older normal hearing with tinnitus, oHI_CO = older hearing impaired control, oHI_T = older hearing impaired with tinnitus.

